# Protein Tyrosine Phosphatase 1B Regulates MicroRNA-208b-Argonaute 2 Association and Thyroid Hormone Responsiveness in Cardiac Hypertrophy

**DOI:** 10.1101/763953

**Authors:** Gérald Coulis, Yanfen Shi, David P. Labbé, Alexandre Bergeron, Fatiha Sahmi, Valérie Vinette, Gérard Karsenty, Bruce G. Allen, Michel L. Tremblay, Jean-Claude Tardif, Benoit Boivin

## Abstract

Elevated reactive oxygen species (ROS) production plays an important role in the pathogenesis of several diseases, including cardiac hypertrophy. While the regulation of diverse sources of ROS is well characterized in the heart, the redox-sensitive targets that contribute to redox signaling remain largely undefined. We now report that protein tyrosine phosphatase 1B (PTP1B) is reversibly oxidized and inactivated in hearts undergoing hypertrophy and that gene deletion of PTP1B in mouse hearts cause an hypertrophic phenotype that is critically exacerbated in mice subjected to pressure overload. Furthermore, we show that PTP1B dephosphorylates Tyr^393^ on argonaute 2, a key component of the RNA-induced silencing complex, and sustains gene silencing in the heart. Our results indicate that PTP1B inactivation and argonaute 2 Tyr^393^ phosphorylation specifically prevents argonaute 2 from interacting with miR-208b. Phosphorylation and inactivation of argonaute 2 in PTP1B cKO mice revealed a mechanism by which defective miR-208b-mediated repression of thyroid hormone receptor-associated protein 1 (THRAP1/MED13) contributes to thyroid hormone-mediated cardiac hypertrophy. In support of this conclusion, inhibiting the synthesis of triiodothyronine (T3), using propylthiouracil, rescued TAC-induced hypertrophy and improved myocardial contractility and systolic function in PTP1B cKO mice. Together, our data illustrate that PTP1B activity exerts a cardioprotective effect in the heart and that redox signaling is tightly linked to thyroid hormone responsiveness and to microRNA-mediated gene silencing in pathological hypertrophy.

## INTRODUCTION

The heart undergoes pathological hypertrophic remodeling in response to chronic stresses such as pressure overload and neurohumoral stimulation (*1*). These pathophysiological stimuli initially trigger a series of electrophysiological, transcriptional, translational, and signaling changes that contribute to temporary adaptations, including cardiomyocyte growth. Sustained exposure, however, is associated with decompensation, dilated cardiomyopathy, arrhythmias, heart failure and ultimately, death (*2, 3*). Previous studies have revealed that reactive oxygen species (ROS) are intimately involved in cardiac hypertrophy, ischemia and heart failure (*4–6*). Indeed, decreasing levels of ROS, either genetically or pharmacologically prevents the transition from cardiac hypertrophy to heart failure (*7–9*) and protects the heart from ischemia-reperfusion injury (*10*). In addition, transgenic mice overexpressing a master gene of the cellular anti-oxidant response, NRF2 (nuclear factor erythroid-2 related factor) are protected from pathological hypertrophy caused by hemodynamic stress (*11*) and mice ectopically expressing a tunable hydrogen peroxide-generating oxidase in the heart display significant systolic dysfunction and hypertrophy (*12*). Nonetheless, despite the convincing causal relationship between increased ROS levels and the pathogenesis of cardiovascular diseases, most redox-regulated signaling events that promote cardiac hypertrophy have yet to be elucidated.

Regulation of protein phosphorylation by redox signaling has emerged as a critical determinant of normal *versus* pathological signaling (*13–15*). In a physiological context, reversible oxidation of reactive cysteine residues within proteins elicit a spectrum of structural alterations that allow ROS production to be coupled to changes in protein activity and cell function (*15*). The large superfamily of cysteine-dependent protein tyrosine phosphatases (PTPs) integrate redox-modifications to regulation of protein phosphorylation (*16–20*). PTPs possess a catalytic center with a unique architecture that maintains the side-chain of their conserved catalytic cysteine residue in a reactive thiolate form. This structural characteristic grants PTPs with the ability to catalyze phosphoryl hydrolysis of their respective substrates and allows the dynamic cellular redox microenvironment to fine-tune dephosphorylation by inhibiting the catalytic activity of PTPs through reversible oxidation of their catalytic cysteine residue (*15, 16, 20*). To date, no oxidized PTPs have been identified in the progression of cardiac hypertrophy and heart failure.

Few members of the PTP superfamily have been studied in the heart. Among them, mitogen-activated protein kinase phosphatases (MKPs) and src-homology 2 domain-containing phosphatase (SHP2) regulate the mitogen-activated protein kinase (MAPK) pathways (i.e. ERKs, JNKs and p38 MAPKs) and their genetic inactivation contributes to cardiac growth and remodeling (*21–23*). In contrast, global deletion of protein tyrosine phosphatase 1B (PTP1B, encoded by the PTPN1 gene) or systemic pharmacological inhibition of the phosphatase in mice confers cardioprotective effects by an unknown mechanism (*23, 24*). Given that studies using tissue-specific PTP1B knockout (KO) mice have revealed PTP1B to control complex organ- and cell-type specific signaling events (*25–30*), we enquired whether cardioprotection observed in the whole-body KO was conferred by cardiomyocyte-targeted PTP1B inactivation.

In testing this hypothesis, we identified PTP1B as a target of ROS signaling in pressure overload-induced hypertrophy. To better understand how PTP1B inactivation contributes to cardiac hypertrophy, we generated a cardiomyocyte-specific PTP1B KO (PTP1B cKO) mouse and we showed that disruption of PTP1B function in cardiomyocytes exacerbated pressure overload-mediated pathological remodeling. We observed that inactivation of PTP1B increased phosphorylation of its substrate, argonaute 2 (AGO2) at Tyr^393^, a phosphorylation event that prevents microRNA (miRNA) loading onto AGO2 (*31*). Herein, we show that loading of miR-208b onto AGO2 is compromised when PTP1B is inactivated, leading to defective miRNA-mediated repression of thyroid hormone receptor-associated protein 1 (THRAP1, also known as MED13) and triiodothyronine (T3)-mediated hypertrophy. Our findings shed light on the role of PTP1B inactivation by ROS in the regulation of gene silencing and the thyroid hormone response in pathological cardiac hypertrophy.

## RESULTS

### Generation of cardiomyocyte-specific PTP1B knockout mice

The function of PTP1B is unknown in the heart. To gain some insight into the cardiac function of PTP1B, we first assessed whether PTP1B was reversibly oxidized and inactivated in hearts undergoing hypertrophy. We measured the reversible oxidation of PTP1B in the heart using a modified cysteinyl-labeling assay, a 3-step method where reversible oxidation of the invariant catalytic cysteine residue of PTPs is specifically converted to a biotin modification, which allows high-affinity purification using streptavidin (Figure 1A) (*17, 18*). Immunoblotting the biotin-streptavidin complexes, generated using the cysteinyl-labeling assay in hearts from sham-operated (control) and mice one or two weeks after constriction of the transverse aorta (TAC) to induce pressure overload and subsequent hypertrophy, revealed that PTP1B was reversibly oxidized in hearts at 1 and 2 weeks post-TAC (Figure 1B). In contrast, the level of total PTP1B was unaffected by TAC. This demonstrates for the first time that PTP1B is inactivated by reversible oxidation in hearts undergoing hypertrophy. Since knocking out a PTP gene effectively reproduces ROS-mediated PTP inactivation and increases the phosphorylation of its substrates, we generated a line of cardiomyocyte-specific PTP1B knockout (PTP1B cKO) mice to understand how PTP1B inactivation in cardiomyocytes affects the development of hypertrophy. Cardiac-specific deletion of PTP1B was achieved by crossing mice with *loxP* sites flanking exons 6 to 8 of PTP1B (the catalytic domain is contained in exons 7 and 8) (*30*) with a line of α-MHC-Cre mice previously validated for their lack of cardiotoxicity (Figure 1C) (*32*). Genetic background, exon deletion, PTP1B mRNA levels and protein expression were validated by PCR and western blotting, respectively (Figure 1D-F). In Figure 1D, PCR genotyping of PTP1B^WT/F^, PTP1B^F/F^ and α-MHC-Cre/PTP1B^F/F^ (PTP1B cKO) mice revealed a two-band pattern corresponding to heterozygotes possessing one WT (230 bp) and one floxed allele (280 bp). A single band at 280 bp was observed for PTP1B^F/F^ mice also possessing the α-MHC-Cre mice allele. PCR of PTP1B mRNA from hearts, lungs and livers of α-MHC-Cre/PTP1B^F/F^ mice confirmed that PTP1B was effectively and specifically knocked out in cardiac tissue (Figure 2E), as confirmed by PTP1B immunoblots of adult cardiomyocyte lysates from PTP1B^F/F^ or PTP1B cKO mice (Figure 1F).

**Figure 1.**
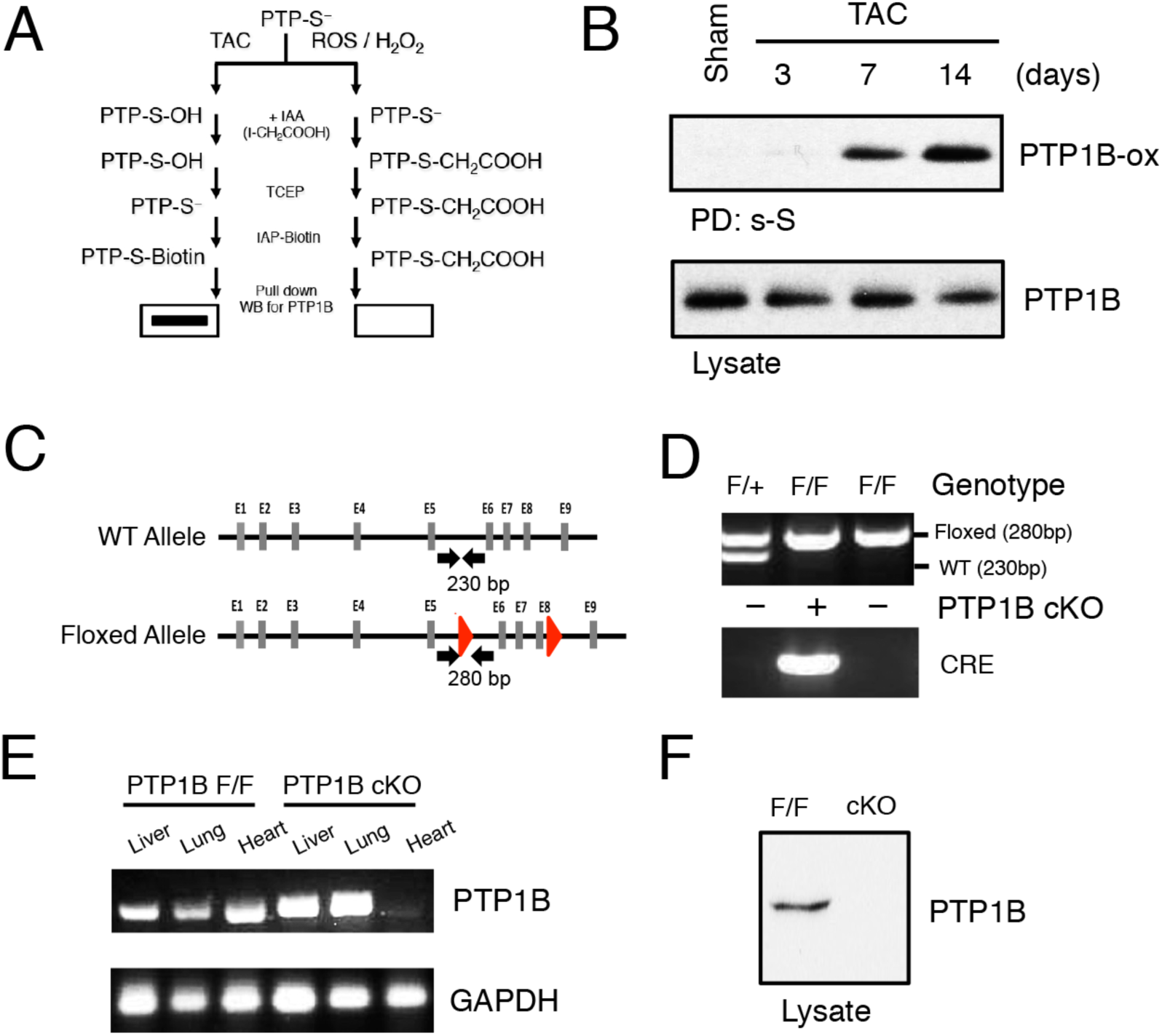
Generation of cardiomyocyte-specific PTP1B knockout mice. A) Schematic outline of the cysteinyl-labeling assay. B) Sham or TAC hearts were subjected to the cysteinyl-labeling assay, using biotinylated IAP at pH 5.5. Proteins enriched in the cysteinyl-labeling assay were resolved by SDS-PAGE and immunoblotted using an anti-PTP1B antibody (upper panel). Total PTP1B immunoreactivity was measured in 5% of the lysate that was used in the cysteinyl-labeling assay. C) Domain structure of PTP1B. Two *loxP* sites were added to flank exons 6 to 8 since the catalytic domain of PTP1B is contained in exons 7 and 8. Small arrows indicate locations targeted by primers used for genotyping. D) PCR genotyping of mice generated by breeding PTP1B^F/F^ with αMHC-Cre/PTP1B^F/F^ mice. A two-band pattern corresponded to heterozygotes possessing one WT (230 bp) and one floxed allele (280 bp) and a single band at 280 bp corresponded to PTP1B^F/F^ mice. E) PTP1B mRNA from hearts, lungs and livers of PTP1B^F/F^ and αMHC-Cre/PTP1B^F/F^ (PTP1B cKO) mice were measured by PCR. GAPDH levels were measured to assess RNA input. F) Lysates from adult cardiomyocytes, freshly isolated from PTP1B^F/F^ or PTP1B cKO mice were resolved by SDS-PAGE and immunoblotted using an anti-PTP1B antibody.

**Figure 2.**
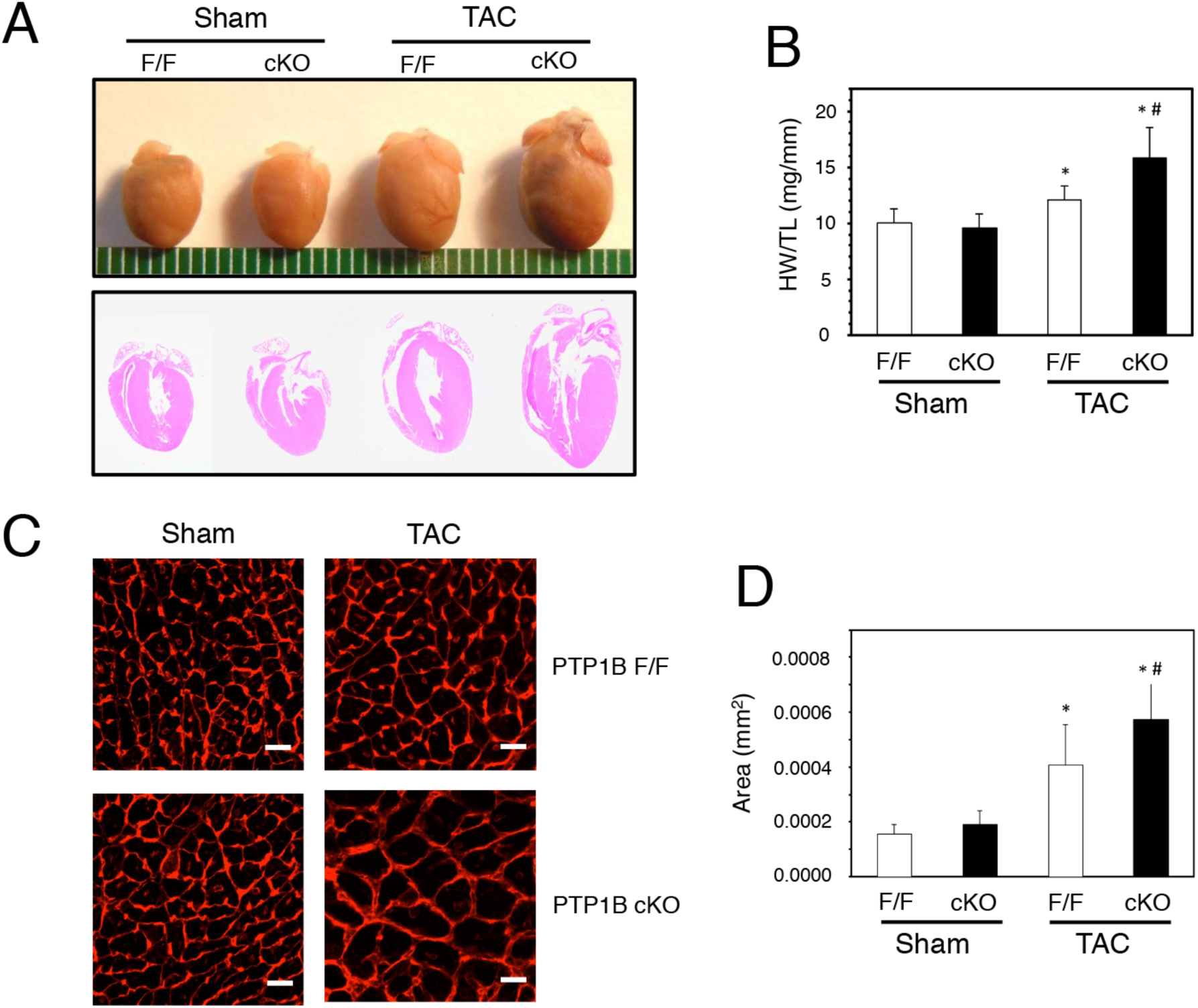
Deletion of PTP1B in cardiomyocytes exacerbates development of cardiac hypertrophy. A) Anatomical view and histological hematoxylin and eosin (HE)-stained sections from hearts from sham and TAC PTP1B^F/F^ and PTP1B cKO mice 4 weeks post-surgery. B) Quantitative analysis of heart weight, normalized to tibia length (HW/TL), for hearts from sham and TAC PTP1B^F/F^ and PTP1B cKO mice 4 weeks post-surgery. C) Representative left ventricular cross-sectional sections from hearts from sham and TAC PTP1B^F/F^ and PTP1B cKO mice 4 weeks post-surgery labeled with Texas Red-X conjugated wheat germ agglutinin (WGA). *Scale bars* = 50 μm D) Quantitative left ventricular cardiac myocyte cross-sectional area from 2000-3000 WGA-labeled cardiomyocytes from sham and TAC PTP1B^F/F^ and PTP1B cKO mice 4 weeks post-surgery. Data represented as mean ± standard error of the mean (SEM). Statistical analyses were done by Student’s t-test. * *P* < 0.05 vs. corresponding Sham group value; # *P* < 0.05 vs. TAC PTP1B^F/F^ group value.

### Cardiomyocyte-specific deletion of PTP1B exacerbates hypertrophy

As anticipated based on previous studies using global PTP1B KO mice, PTP1B cKO mice were healthy and showed no apparent phenotype under basal conditions (*25*). However, echocardiographic imaging of 6-8 week-old PTP1B cKO mice revealed differences between PTP1B^F/F^ and PTP1B cKO (Table S1). Specifically, PTP1B cKO mice showed small but significant bradycardia (HR), left ventricular enlargement (LVDd), increased left ventricular mass (LVM) and decreased left ventricular systolic (FS) and diastolic function (IVRT). To test whether inactivation of PTP1B directly contributes to pathological cardiac hypertrophy, we subjected control PTP1B^F/F^ and PTP1B cKO mice to TAC. As expected, 4 weeks post-surgery, control TAC PTP1B^F/F^ mice had increased heart size and left ventricular wall thickness (Figure 2A, Sham *versus* TAC F/F). However, these morphological changes were significantly increased in TAC PTP1B cKO mice (Figure 2A, Sham *versus* TAC cKO) and gravimetric measurements confirmed that TAC PTP1B cKO mice had a significant increase in heart weight to tibia length ratio (HW/TL) (Figure 2B) relative to TAC PTP1B^F/F^ mice. Furthermore, the increased heart weight of PTP1B cKO mice correlated with a quantitative increase in cardiomyocyte cross-sectional area, an index of cardiomyocyte size (Figure 2C, D). Collectively, these data demonstrate that PTP1B inactivation in cardiomyocytes promote cardiac hypertrophy in vivo.

### PTP1B exerts a cardioprotective function in response to pressure overload

To evaluate cardiac function in PTP1B cKO mice, echocardiographic imaging was performed. Examining mice 4 weeks post-TAC revealed that the left ventricular filling patterns of PTP1B cKO hearts mice were significantly altered (Figure 3A). Left ventricular systolic function was significantly reduced, as indicated by decreased fractional shortening (FS) and ejection fraction (EF). In addition, left atrial diameter (LAD) was increased, indicating left ventricular diastolic dysfunction (Figure 3B-D and Table S2). At the molecular level, cardiac hypertrophy is characterized by increased expression of fetal genes such as β-MHC (*Myh7*), ANP (*Nppa*) and BNP (*Nppb*) in the adult heart (*33, 34*). Consistent with our morphometric and hemodynamic observations, qPCR analysis revealed that TAC significantly increased β-MHC, ANP and BNP mRNA levels in control PTP1B^F/F^ mice (Figure 3E). Surprisingly, whereas ANP and BNP mRNA were also significantly higher in PTP1B cKO mice subjected to TAC, the expected increase in β-MHC transcript levels leading to the switch in MHC isoform expression observed in control TAC PTP1B^F/F^ mice was compromised in TAC PTP1B cKO littermates. Taken together, these data indicate that loss of PTP1B function exacerbates both hypertrophy and the impairment of cardiac function induced by pressure overload.

**Figure 3.**
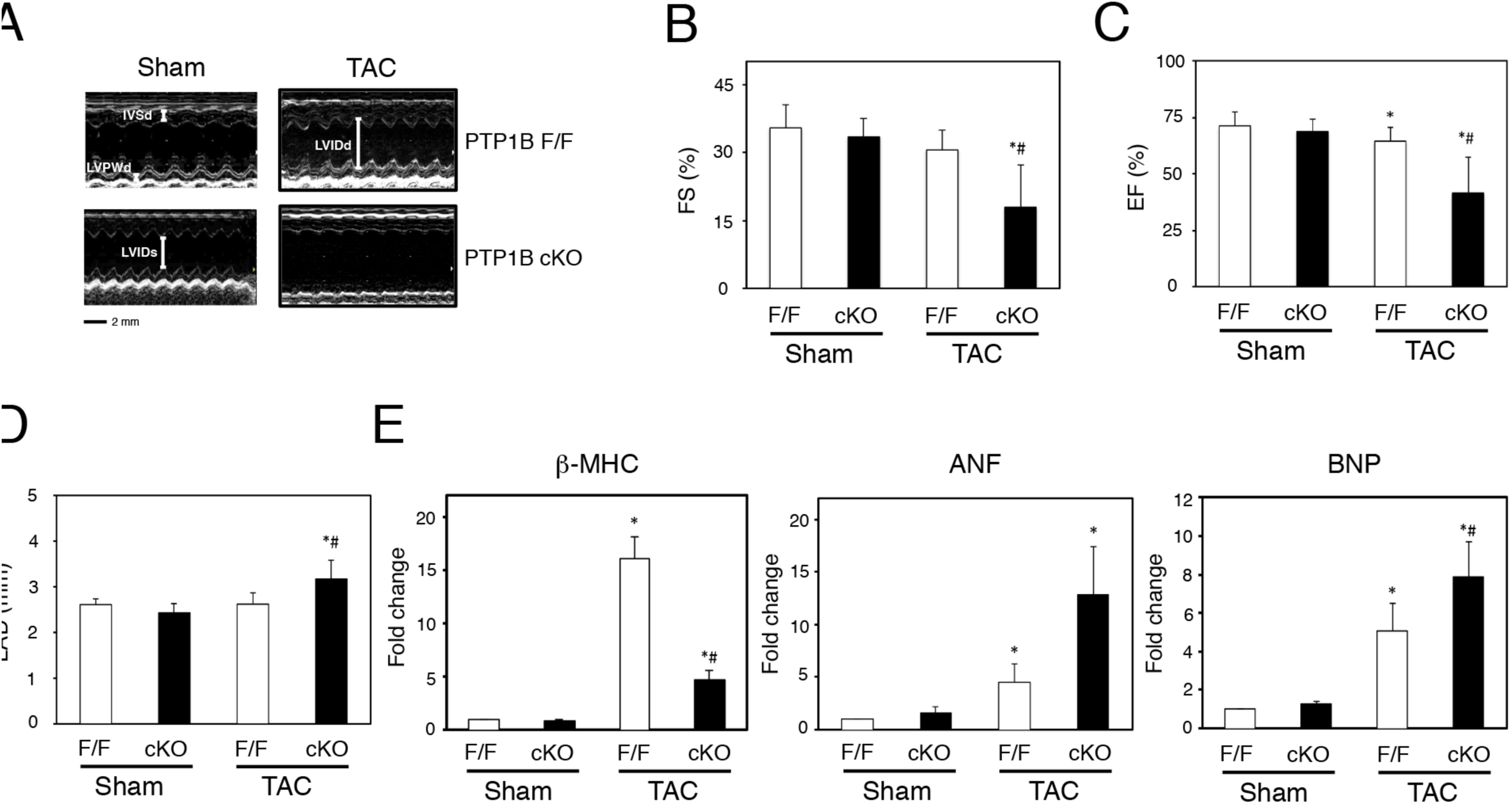
Cardiomyocyte-specific deletion of PTP1B impairs left ventricular performance and β-MHC gene expression. A) Representative M-mode echocardiograms of sham and TAC PTP1B^F/F^ and PTP1B cKO mice 4 weeks post-surgery. The IVSd (interventricular septal end diastole) and LVPWd (Left ventricular posterior wall end diastole) dimensions were used to determine left ventricular hypertrophy, LVIDd and LVIDs (left ventricular internal diameter end diastole and end systole, respectively) were used to calculate fractional shortening (FS). B) Changes in left ventricular fractional shortening, C) left ventricular ejection fraction, and D) isovolumic relaxation time (IVRT) in sham and TAC PTP1B^F/F^ and PTP1B cKO mice 4 weeks post-surgery. E) Quantitation of molecular indicators of hypertrophy β-MHC/GAPDH, ANF/GAPDH and BNP/GAPDH mRNA by qRT-PCR. Values are presented as mean ± SEM for 3 or more independent experiments. * *P* < 0.0001 vs. respective sham-operated groups; # *P* < 0.0001 vs. TAC PTP1B^F/F^ group.

### PTP1B regulates the association between miRNA-208b and AGO2 in cardiac hypertrophy

MHC isoforms alpha and beta differ in their ability to generate velocity of shortening (*35*) and the switch from the faster α-­-MHC to the slower β-MHC isoform that occurs in cardiac hypertrophy is generally thought to reduce contractile performance and to contribute to functional maladaptation (*36–38*). Since exacerbated hypertrophy is observed in absence of β-MHC expression in PTP1B cKO mice, we reasoned that understanding how PTP1B inactivation prevents the expression of β-MHC would shed some light on the molecular pathways underlying the PTP1B cKO hypertrophic phenotype. The MHC switch is regulated by miRNAs and thyroid hormones in development and hypertrophy (*38–40*). Hence, we first tested whether miRNA populations were affected by PTP1B inactivation and TAC. Four weeks post-surgery, total RNA from TAC and sham PTP1B^F/F^ and PTP1B cKO hearts was isolated and hybridized to microarrays that report the expression status of 1908 murine miRNAs. Cardiac hypertrophy resulted in induction and repression of miRNAs that partially overlapped in PTP1B^F/F^ or PTP1B cKO mice (Figure S1). We identified miR-208b (identified by * in Figure S1), a cardiac-restricted miRNA (myomiR) known to regulate the MHC switch, among the top 20 miRNAs showing most change in expression in PTP1B^F/F^ and PTP1B cKO subsets 4 weeks post-TAC (i.e. log_2_ values relative to sham-operated hearts) (*39*). MiR-208a and miR-208b, respectively encoded by the myosin genes *Myh6* (α-MHC) and *Myh7* (β-MHC), mediate actions of thyroid hormone in the heart (*40*). Although they possess the same seed sequence and are thought to silence an overlapping selection of transcripts in the heart (*39*), their respective regulation in cardiac hypertrophy is unclear. Quantitative PCR confirmed that expression of miR-208b but not miR-208a increased in cardiac hypertrophy (Figure 4A, B). We recently reported that PTP1B activity contributes to maintaining functional miRNA-loaded AGO2, the spearhead of the RNA-induced silencing complex (RISC) gene silencing machinery (*31, 41*). Our results indicate that TAC-induced β-MHC transcript expression was compromised in PTP1B cKO hearts (Figure 3D). To test if the difference in β-MHC expression in PTP1B cKO hearts resulted from decreased silencing by miR-208b, we investigated whether the association between miR-208 isoforms and AGO2 was affected in sham and TAC-treated PTP1B^F/F^ and cKO hearts. Quantitative PCR revealed that miR-208a association with AGO2 was unaffected by PTP1B KO or following TAC surgery (Figure 4C). In contrast, the association between AGO2 and miR-208b increased approximately 1000-fold in hypertrophic TAC PTP1B^F/F^ hearts; however, miR-208b did not associate with AGO2 in TAC PTP1B cKO hearts, despite increased expression of the miRNA (Figure 4D). This suggests that PTP1B inactivation uncouples the expression of specific miRNAs from their ability to post-transcriptionally repress genes in the heart.

**Figure 4.**
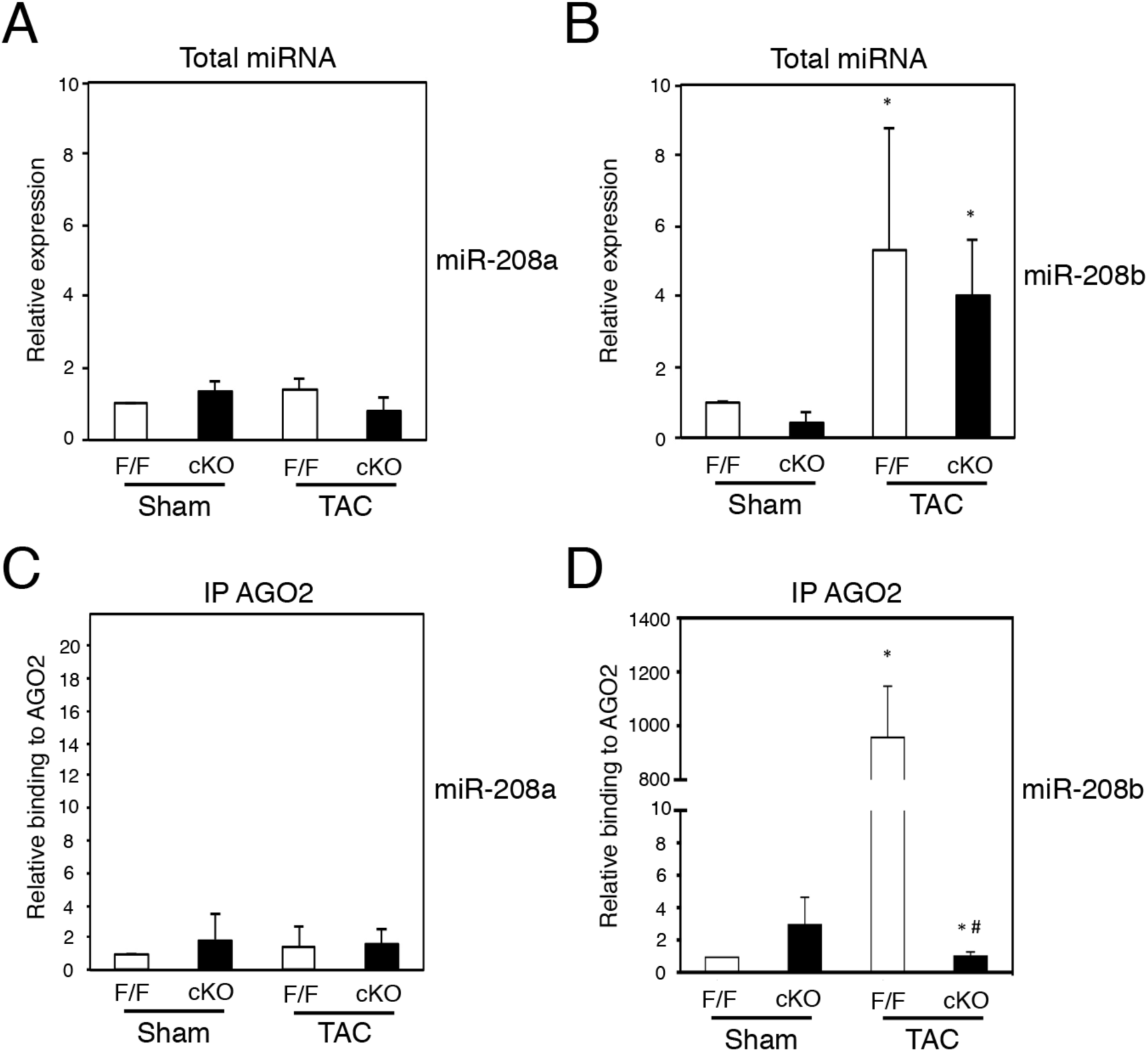
PTP1B regulates the association of miR-208b with AGO2 in hypertrophic hearts. A) and B) Quantitation of miR-208a and miR-208b levels by qRT-PCR in hearts from sham and TAC PTP1B^F/F^ and PTP1B cKO mice 4 weeks post-surgery. C and D) AGO2 was immunoprecipitated from heart lysates from sham and TAC PTP1B^F/F^ and PTP1B cKO mice 4 weeks post-surgery, and AGO2-associated miRNAs were extracted. After cDNAs were synthesized, AGO2-bound miR-208a and miR-208b were assessed by qRT-PCR. MiRNA expression levels and AGO2-enrichment was normalized to 1 in sham PTP1B^F/F^ hearts. Representative data are presented as mean ± SEM for 3 or more independent experiments. * *P* < 0.05 vs. respective sham-operated groups; # *P* < 0.05 vs. TAC PTP1B^F/F^ group.

### AGO2 is a substrate of PTP1B in cardiomyocytes and in hearts undergoing hypertrophy

The importance of miRNAs in cardiovascular development and disease is well established (*39, 42, 43*); however, AGO2 regulation has not been examined in this context. PTP1B regulates gene silencing through dephosphorylation of AGO2 at Tyr^393^ (*31*), a phosphorylation event that impairs miRNA loading onto AGO2 in cells (*31, 44*). We therefore tested if inactivation of PTP1B contributes to AGO2 phosphorylation and decreased association with miR-208b in cardiomyocytes. We first expressed a substrate-trapping mutant PTP1B D181A (DA) (*45*) in cardiomyocytes in which an activated *RAS* allele (*H-RAS^V12^*) was introduced by a retroviral vector for a 2-, 3- or 4-day period in order to induce hypertrophy (*46*). Immunoprecipitation of PTP1B (DA) from cellular extracts of cardiomyocytes expressing H-RAS^V12^ confirmed that AGO2 was part of the PTP1B (DA) immune complex and a substrate of the phosphatase in hypertrophic cardiomyocytes (Figure 5A). To further assess whether AGO2 is a substrate for PTP1B in the hypertrophying myocardium, we generated a Cre/loxP conditional *Rosa26*-targeted transgenic mouse that overexpresses a 3X-Flag-*PTPN1* (DA) transgene together with an eGFP/luciferase reporter (Figure S2A). PTP1B (DA) expression was achieved by crossing these mice to *Meox2*-Cre mice. Excision of the STOP cassette was assessed by live imaging as previously described (Figure S2B) (*47*). Two weeks post TAC, Flag-PTP1B immunoprecipitation from heart lysates validated PTP1B (DA) expression in *Rosa26^PTPN1-DA/WT^* mice compared to *Rosa26^WT/WT^* control littermates and revealed that AGO2 was a substrate of PTP1B in vivo (Figure 5B). The prolonged association between AGO2 and the PTP1B substrate-trapping mutant confirms that AGO2 is a substrate of PTP1B in cardiomyocytes and in the heart.

**Figure 5.**
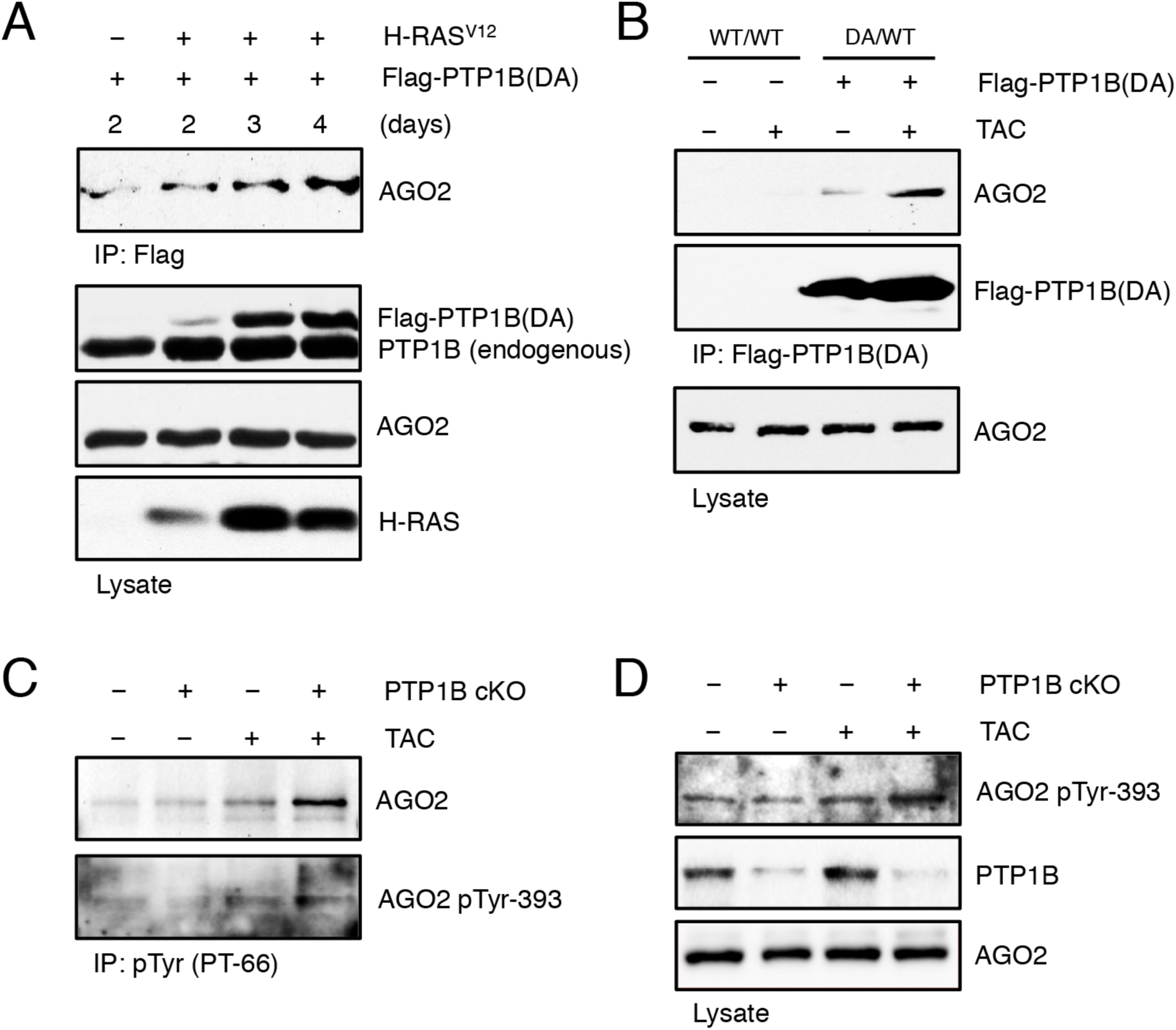
PTP1B dephosphorylates Argonaute 2 at Tyr^393^ in hypertrophic hearts. A) Lysates from cardiomyocytes expressing Flag-PTP1B (DA) or H-RAS^V12^ and Flag-PTP1B (DA) for 2, 3 or 4 days were subjected to substrate-trapping and immunoblotted with an anti-AGO2 antibody. Lysates were resolved by SDS-PAGE and probed for total PTP1B, AGO2 and H-RAS immunoreactivity to confirm the expression of the mutant proteins and AGO2. B) Lysates from sham or TAC hearts from control transgenic mice (*Rosa26^WT/WT^*) or from transgenic mice expressing Flag-PTP1B DA (*Rosa26^PTPN1-DA/WT^*), two weeks post-surgery, were subjected to substrate-trapping and blotted with an anti-AGO2 antibody. The efficiency of the immunoprecipitation was monitored by immunoblotting for Flag and AGO2 levels were also monitored in the heart lysates to monitor changes in expression. C) Tyrosine phosphorylated AGO2 was immunoprecipitated from sham and TAC PTP1B^F/F^ and PTP1B cKO mice 4 weeks post-surgery using PT-66. Proteins were resolved by SDS-PAGE, transferred, and immunoblotted using an anti-AGO2 antibody. The membrane was stripped and reblotted for phospho-Tyr^393^ AGO2. D) AGO2 Tyr^393^ phosphorylation was monitored in heart lysates form sham and TAC PTP1B^F/F^ and PTP1B cKO mice 4 weeks post-surgery. Proteins were resolved by SDS-PAGE, transferred, and immunoblotted using an anti-phospho-Tyr^393^ AGO2 antibody, and anti-AGO2 and PTP1B antibodies to illustrate expression levels in PTP1B cKO mice.

We next asked whether PTP1B inhibition increase AGO2 phosphorylation during hypertrophy. Tyrosine-phosphorylated proteins were immunoprecipitated from TAC and sham heart lysates from control or PTP1B cKO mice four weeks post-surgery and AGO2 was visualized by immunoblot. As expected, AGO2 was tyrosine phosphorylated in TAC PTP1B^F/F^ hearts and hyperphosphorylated in TAC PTP1B cKO hearts (Figure 5C, upper panel). Similarly, phosphorylation of AGO2 at Tyr^393^ was also more pronounced in PT-66 (anti-phosphotyrosine) immunoprecipitates (Figure 5C, lower panel) and heart lysates of TAC PTP1B cKO mice, as compared to TAC PTP1B^F/F^ littermates and sham-operated control mice (Figure 5D). Consistent with PTP1B oxidation in cardiac hypertrophy previously observed in figure 1, AGO2 was phosphorylated at Tyr^393^ in heart lysates from mice one and two weeks post-TAC (Figure S3). This indicates that PTP1B dephosphorylates AGO2 in hearts and that, in response to pressure overload, reversible oxidation of PTP1B leads to phosphorylation and inactivation of a population of AGO2 in cardiac hypertrophy.

### PTP1B inactivation impairs translational repression of THRAP1

High target-prediction using web-databases (*48, 49*) combined with validated studies (*50, 51*) indicated that miR-208 isoforms target sites within the 3’UTR of THRAP1/MED13 transcripts, a component of the Mediator complex known to repress β-MHC transcription in a thyroid hormone-dependent manner (*39, 40*). This prompted us to assess THRAP1 expression in PTP1B cKO mice subjected to TAC. Since Tyr^393^-phosphorylation effectively prevents AGO2 from interacting with and repressing mRNA targets (*31*), we investigated whether AGO2 inactivation could affect its association with THRAP1 transcripts. AGO2 was immunoprecipitated from sham or TAC PTP1B^F/F^ and PTP1B cKO hearts and AGO2-associated THRAP1 mRNA levels were analyzed by qPCR. The association between AGO2 and THRAP1 mRNA was minimal in sham-operated hearts and increased approximately 4-fold in TAC PTP1B^F/F^ hearts when AGO2 is minimally phosphorylated (Figure 6A). In contrast, the association between AGO2 and THRAP1 mRNA did not increase significantly in TAC PTP1B cKO hearts when AGO2 is phosphorylated and does not interact with miR-208b as shown in Figure 4D. This suggests that phosphorylation of AGO2 on Tyr^393^ decreases its ability to repress THRAP1. Accordingly, while cellular THRAP1 mRNA levels were not significantly affected by TAC (Figure 6B), protein expression of THRAP1 was markedly increased when THRAP1 mRNA is not repressed by AGO2 in TAC PTP1B cKO hearts (Figure 6C). This illustrates that phosphorylation of AGO2 at Tyr^393^ impairs the translational repression of THRAP1 mRNA in hypertrophic hearts, despite the abundance of miR-208 isoforms. As TRβ1 associates with THRAP1 to repress β-MHC (*50, 51*), we then assessed TRβ1 expression in the heart. TRβ1 levels were elevated in control and hypertrophic PTP1B cKO hearts compared to TAC PTP1B^F/F^ mice supporting the idea that PTP1B expression affects thyroid hormone responsiveness in the heart.

**Figure 6.**
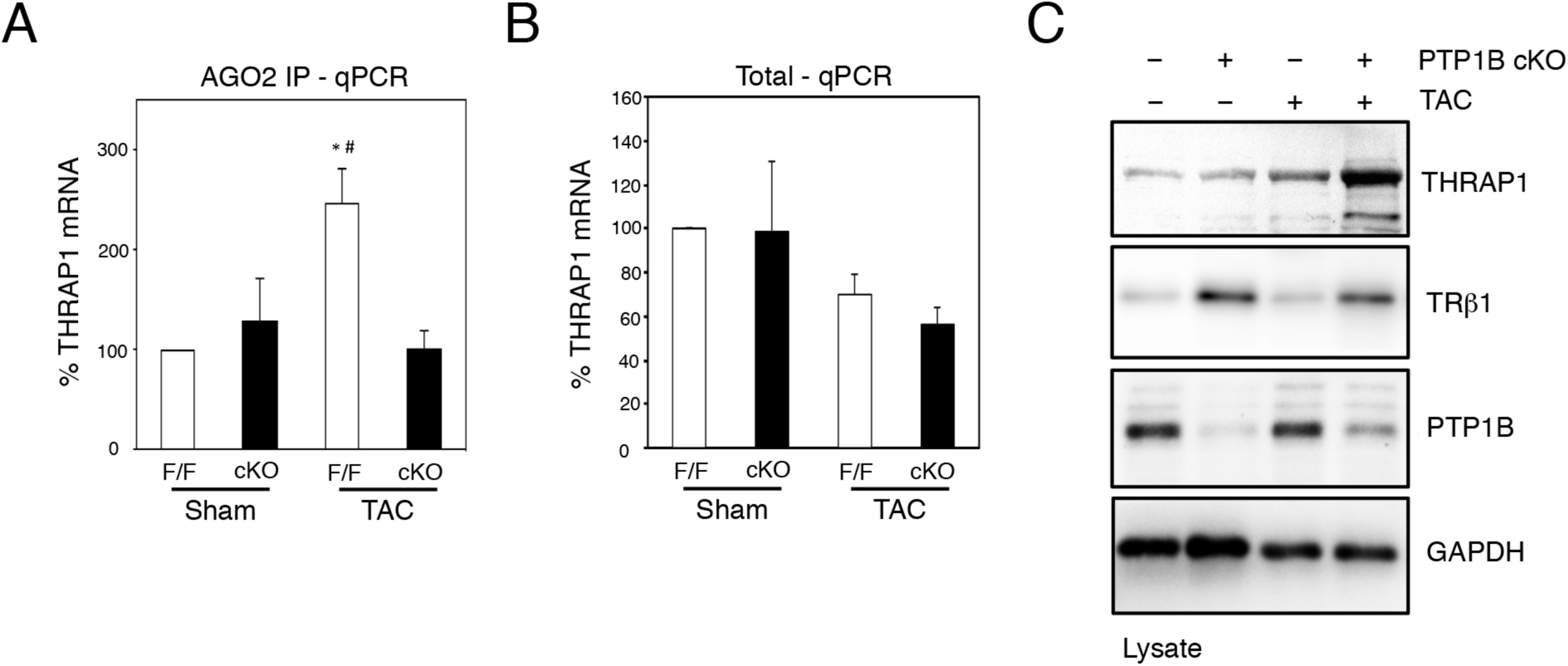
Argonaute 2 inactivation facilitates THRAP1 expression in hypertrophic hearts from PTP1B cKO mice. A) AGO2 was immunoprecipitated from lysates of hearts from sham and TAC PTP1B^F/F^ and PTP1B cKO mice 4 weeks post-surgery. AGO2-associated mRNAs were extracted, cDNAs were synthesized and used for qRT-PCR for THRAP1. mRNA enrichment levels were normalized to 100% in sham PTP1B^F/F^ hearts. B) Quantitation of THRAP1 levels by qRT-PCR in hearts from sham and TAC PTP1B^F/F^ and PTP1B cKO mice 4 weeks post-surgery. THRAP1 mRNA levels were normalized to 100% in sham PTP1B^F/F^ hearts. C) Heart lysates from sham and TAC PTP1B^F/F^ and PTP1B cKO mice 4 weeks post-surgery were resolved by SDS-PAGE, transferred and blotted for THRAP1, PTP1B and GAPDH. Values are presented as mean ± SEM for 3 or more independent experiments.

### Inhibition of T3 synthesis protects PTP1B cKO hearts from pressure-overload

Thyroid hormone signaling, through enhancer and repressor cis-acting elements, respectively located in the promotors of α-MHC and β-MHC genes, plays a critical role in the MHC isoform switch that occurs in hypertrophy and heart failure (*38*). In this context, THRAP1 and TRβ1 co-repress β-MHC expression in a T3-dependent manner (*50, 51*). We examined whether thyroid signaling contributed to impaired myosin switch, exacerbated hypertrophy and cardiac dysfunction observed in PTP1B cKO mice subjected to pressure overload. To test this hypothesis, we fed propylthiouracil (PTU)-containing chow to PTP1B cKO and control mice to inhibit triiodothyronine (T3) synthesis, block T3 signaling, and thus induce hypothyroidism for 4 weeks post-surgery. Gravimetric measurements revealed that blunting thyroid signaling essentially prevented the TAC-induced increase in heart weight to tibia length ratio (HW/TL) in PTP1B cKO mice (Figure 7A). In addition, echocardiographic imaging 4 weeks post-surgery revealed that the left ventricular filling patterns of sham and TAC hearts from PTU-treated mice appeared minimally altered (Figure S4) when compared to the filling patterns of non-PTU treated animals (Figure 3A). Detailed analysis confirmed that TAC PTP1B cKO hearts were protected from systolic dysfunction as indicated from preserved fractional shortening and ejection fraction values (Figure 7B, C). Likewise, diastolic function [as indicated by left atrial diameter (LAD) and heart rate-corrected isovolumic relaxation time (IVRTc)] were also preserved in PTU-treated TAC hearts (Table S3). These observations suggests that preventing thyroid hormone signaling maintains systolic and diastolic function and prevents the development of cardiac hypertrophy when PTP1B is inactivated in response to pressure overload.

**Figure 7.**
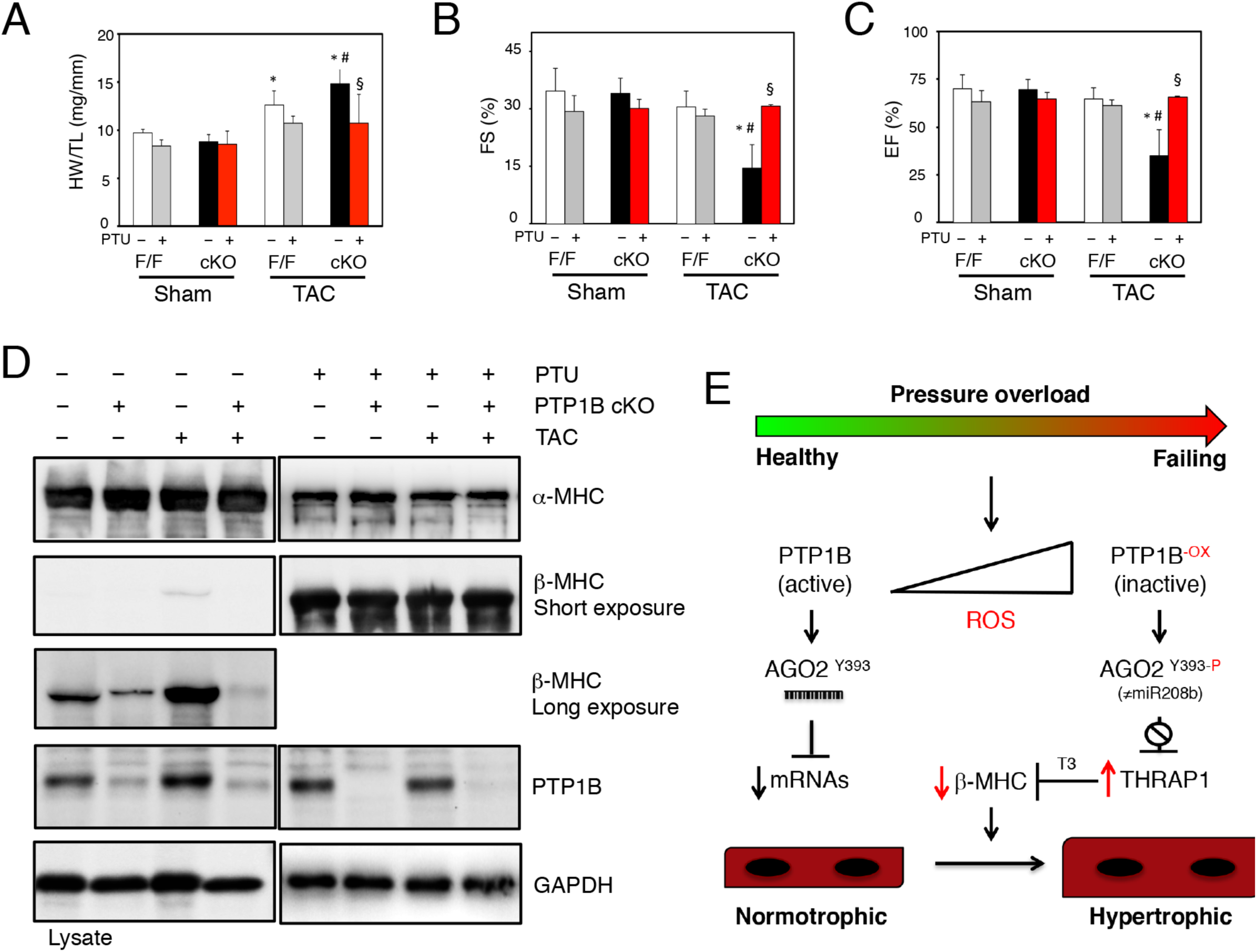
PTP1B regulates thyroid hormone responsiveness of the β-MHC gene and cardiac hypertrophy. A) Quantitative analysis of heart weight, normalized to tibia length (HW/TL), for sham and TAC PTP1B^F/F^ and PTP1B cKO mice 4 weeks post-surgery, treated or not with PTU. B) Echocardiographic analysis of left ventricular fractional shortening (FS) and C) ejection fraction (EF) in sham and TAC PTP1B^F/F^ and PTP1B cKO mice 4 weeks post-surgery, treated or not with PTU. D) Western blot analysis of α-MHC and β-MHC expression in hearts from PTP1B^F/F^ and PTP1B cKO mice at baseline with or without 4 weeks of TAC, treated or not with PTU. E) Model of redox-regulation of PTP1B, AGO2 and thyroid hormone responsiveness in cardiac hypertrophy. Data represented as mean ± SEM. Statistical analyses were done by two-way ANOVA with Bonferonni’s post hoc test for comparisons among multiple groups. * *P* < 0.01 vs. respective sham-operated groups; # *P* < 0.01 vs. TAC PTP1B^F/F^ group; § *P* < 0.01 vs. non-PTU treated TAC PTP1B cKO group.

In order to gain insight into the underlying mechanism of the PTU-induced suppression of the hypertrophic response, we monitored the α-MHC to β-MHC switch as an indicator of T3 signaling in PTU-treated mice subjected to TAC. α-MHC protein levels were similar in all non-PTU-fed mice (Figure 7D, upper left panel). However, preventing T3 synthesis caused a ∼ 2-fold decrease in α-MHC immunoreactivity under all conditions and genotypes (Figure 7D, upper right panel). On the other hand, we could observe that β-MHC immunoreactivity was markedly reduced in both sham and TAC PTP1B cKO mice compared to PTP1B^F/F^ mice (Figure 7D, 3^rd^ panel showing a long exposure). PTU treatment caused a ∼ 30-fold increase in β-MHC immunoreactivity (Figure 7D, 2^nd^ panels showing a short exposure) and a marked increase in mRNA levels (Figure S5) in sham and TAC PTP1B cKO mice, as well as in PTP1B^F/F^ mice. This supports that T3 signaling is blunted and that TRβ1 and THRAP1 do not repress β-MHC transcription in PTP1B cKO hypothyroid animals. Quantification of MHC immunoreactivity indicates that β-MHC represents ∼ 60% of MHC expression in PTU-treated hearts (Figure S6). Thus, increased expression of the slower β-MHC isoform can occur and contribute to contractile performance in hearts from hypothyroid mice subjected to pressure overload. Taken together, our studies support a model in which PTP1B inhibition by ROS regulates AGO2-mediated silencing and modulates thyroid-hormone responsiveness in cardiac hypertrophy (Figure 7E).

## DISCUSSION

Although cardiac redox signaling regulates the activity of proteins and pathways involved in pathological hypertrophy and heart failure, the redox regulation and inactivation of members of the PTP family has not been explored in the heart (*4*). We addressed the cardiac function of PTP1B by monitoring its inactivation by reversible oxidation utilizing a cysteinyl-labeling assay and by generating a line of cardiac-specific PTP1B KO mice. Our results demonstrate that PTP1B inactivation is an integral part of the signaling events leading to pressure overload-induced cardiac hypertrophy. Our data also indicate that reversible oxidation of PTP1B leads to phosphorylation of AGO2, which in turn prevents the association between AGO2 and miR-208b and AGO2-mediated silencing of THRAP1. This mechanism contributes to increased thyroid hormone-mediated cardiac dysfunction and hypertrophy (Figure 7). Together, the results from our study shed new light on the role of PTP1B in regulating thyroid hormone responsiveness in the heart and on how redox signaling is tightly linked to gene silencing and cardiac remodeling.

It is well established that members of the PTP family display substrate specificity in a cell-specific manner (*16*). Gene-targeting studies, using either global PTP1B KO or tissue-specific PTP1B KOs (e.g. muscle, liver, adipose tissue, endothelium, bones, etc.) have demonstrated that PTP1B displays complex cell-specific functions ranging from attenuation of insulin receptor signaling, to enhancing HER2 signaling in breast cancer (*25–30, 52, 53*). Previous studies using either PTP1B inhibitors or a global PTP1B KO mouse, revealed that global gene-inactivation or systemic inhibition of PTP1B decrease cardiac remodeling and dysfunction following myocardial infarction models; however, the important peripheral effects of these approaches, known off-target effects of PTP1B inhibitors, and known cell-specific functions of the phosphatase preclude one from drawing conclusions on the role of PTP1B in heart (Thiebaut et al. 2018). We identified PTP1B as a target of ROS in pressure overload-induced hypertrophy and showed that disruption of PTP1B function is an important checkpoint involved in pathological remodeling in vivo. Our experimental results demonstrate that cardiomyocyte-specific deletion of PTP1B exacerbates pressure-overload-induced hypertrophy, clearly identifying a cardioprotective function for PTP1B in heart failure. PTP1B is widely expressed in the cardiovascular system (*54*) and it will be important to further characterize the role of PTP1B in non-cardiomyocytes to clarify the mechanisms underlying the peripheral cardioprotective function of PTP1B inhibition.

Several lines of genetic evidence indicate that miRNAs play key roles in mediating stress responses and in cardiovascular development and disease (*43*). Genetic deletion and overexpression of miR-208a in mice have shown the importance of myomiRs in regulating MHC isoforms in heart disease (*50, 51, 55*). Knockout of miR-208a, encoded by intron 27 of α-MHC, compromises the expression of β-MHC, and importantly, it also compromises the expression of miR-208b located within intron 31 of the β-MHC gene. Notably, these studies have shown that knocking out miR-208a blunts the hypertrophic response in TAC hearts and that transgenic expression of miR-208a is sufficient to induce hypertrophy (*50, 51*). Since miR-208a levels are unaffected by TAC-induced pressure overload, it would be important to understand how miR-208a contribute to cardiac hypertrophy. Our results stress that preventing AGO2 from interacting with miR-208b promotes pathological cardiac hypertrophy and suggest that part of the signaling attributed to miR-208a gene deletion (*50, 51*) may be caused by the repression of miR-208b in these mice, as mice in which miR-208a is knocked out are effectively miR-208a and miR-208b double KOs (*55*). Our data indicate miR-208b is the only isoform to display increased abundance in cardiac hypertrophy. We show that miR-208b expression is augmented several fold, and most importantly that its association with AGO2 increase a 1000-fold in cardiac hypertrophy, in contrast to the association between miR-208a and AGO2 which is not altered. Revealing different patterns of AGO2 and miR208a/b association, regulated by PTP1B, supports the existence of unique AGO2 complexes in the heart and suggests that further defining miRNA and AGO-specific complexes may yield important insight into post-transcriptional regulation.

Tyr^393^ of AGO2 is located close to the DICER-binding domain, and phosphorylation of this residue impairs both the AGO2-DICER interaction and miRNA loading onto AGO2 in cells (*31, 44, 56, 57*). We recently demonstrated that AGO2 phospho-Tyr^393^ is a direct substrate for PTP1B and showed that reversible oxidation of PTP1B and the resulting AGO2 inactivation could override the miRNA expression program and compromise post-transcriptional regulation of a subset of genes in senescent fibroblasts (*31, 41*). Our data presented herein clearly establishes a function for this pathway in the heart. We show that despite the increased expression of miR-208b, the association between miR-208b and AGO2 is compromised in hypertrophic PTP1B cKO hearts, when AGO2 is phosphorylated on Tyr^393^. This is consistent with the inactivation of a population of AGO2 in hypertrophic PTP1B cKO hearts. Moreover, in our hands, increased AGO2 inactivation in hypertrophic PTP1B cKO hearts prevents miR-208b from targeting and repressing specific mRNAs, including THRAP1/MED13, involved in thyroid hormone signaling and repression of β-MHC transcription. In addition, we also show that TRβ1 expression is increased in PTP1B cKO hearts. Collectively, our data suggest that decreased repression of THRAP1 via AGO2 inactivation is part of a mechanism by which PTP1B inactivation affects thyroid hormone responsiveness during pressure overload-induced hypertrophy; however, increased TRβ1 expression in PTP1B cKO hearts also suggests that regulation of T3-signaling by PTP1B may also occur in other tissues.

The expression of β-MHC and α-MHC isoforms is regulated in a developmental stage-specific and thyroid hormone-dependent manner. In mice, there is a β-MHC to α-MHC transition immediately before birth, with the faster isoform, α-MHC, becoming the major isoform (∼ 90%), whereas in humans and other large mammals the slower β-MHC isoform becomes the predominant isoform in the adult heart (*38*). It is well established, both in rodents and humans, that β-MHC expression increases at the expense of α-MHC in cardiac hypertrophy and hypothyroidism. However its role as a molecular indicator of pathological hypertrophy remains unclear since studies have also reported that cellular hypertrophy is not always accompanied by β-MHC re-expression (*58, 59*), and that β-MHC re-expression occurs only in non-hypertrophic (i.e. normotrophic), α-MHC-expressing-myocytes in pressure-overload hypertrophy (*60*). Our results show that PTP1B cKO mice only weakly express β-MHC after 4 weeks of pressure-overload. These mice presented systolic and diastolic dysfunction, as well as increased left ventricular hypertrophy when compared to control mice. Inhibiting T3 synthesis prevented the pro-hypertrophic phenotype observed in PTP1B cKO mice subjected to pressure-overload, suggesting that T3-mediated hypertrophy occurs downstream of PTP1B. Notably, PTU-treated mice express nearly 30-times the amount of β-MHC compared to non-PTU treated mice without showing a proportional repression of α-MHC expression. Rather, a 2-fold repression of α-MHC expression was observed in PTU-treated mice. Since the absence of α-MHC is deleterious to the heart in mice (*37*) and in humans (*61*), we should perhaps consider that β-MHC expression, known to cause slower but efficient contractility (*38, 62*), can be beneficial to the cardiac function, depending on the expression level of α-MHC in hypertrophic myocardium.

The signaling pathway identified herein reveals that PTP1B is a regulator of thyroid-hormone-mediated cardiac hypertrophy. We show that inactivation of PTP1B by ROS leads to phosphorylation of AGO2 at Tyr^393^ and prevents AGO2 from associating with miR-208b and from post-transcriptionally regulating THRAP1/MED13 mRNA as well as other unidentified transcripts. Our results support the hypothesis that prolonged inactivation of specific PTPs by increased redox signaling is an important factor in cardiac pathology, and contributes to the expanding number of studies characterizing the molecular mechanisms that regulate the activity of miRNA-mediated gene silencing. We anticipate that further characterization of the reversibly oxidized PTPs in the heart will identify additional roles for these enzymes in advanced stages of pathological cardiac hypertrophy and heart failure.

## MATERIALS AND METHODS

### Generation of PTP1B cKO mice

In order to specifically delete PTP1B in cardiomyocytes (PTP1B cKO), C57BL/6 mice harboring two *loxP* sites flanking the catalytic domain of PTP1B at exons 6 and 8 (*30*) were crossed with mice expressing the α-MHC-Cre transgene (Jackson Laboratory). Genotyping was performed by PCR using DNA extraction from tail clips (*63*) and gene-specific primers corresponding to PTP1B lox (forward: 5’-CCTGGACTGAGGCTTTCTAGGC-3’; reverse: 5’-GTCTCTGGTGCTGCTCTGAATTGC-3’) and Cre (forward : 5’-CGTACTGACGGTGGGAGAAT-3’; reverse : 5’-ACCAGGCCAGGTATCACTGA-3’). Cardiac knockout was confirmed by RT-PCR. Total RNA was isolated from various tissues of PTP1B^F/F^ or PTP1B cKO mouse neonates and was subjected to semi-quantitative RT-PCR using SuperScript One-Step (Invitrogen) and primers detecting a 171bp region within PTP1B exon 6 (forward: 5’-CACATGGCCTGACTTTGGAG-3’; reverse : 5’-AGTAAGAGGCAGGTGTCAGC-3’). All mice were housed in a temperature-controlled barrier facility on a 12-hour light-dark circadian cycle with access to standard chow and water. Animal procedures described in this study were approved by the Ethics Committee at the Montreal Heart Institute.

### Generation of PTPN1-DA mice

A Gateway-compatible *Rosa26* locus targeting vector (*64*) was used to generate a *Cre/loxP* conditional *Rosa26*-targeted transgenic mouse that overexpresses human PTP1B D181A (DA) mutant using a strategy described previously (*47*) (Figure S1A). Whole-body PTP1B(DA) expression was achieved by crossing *Rosa26^PTPN1-DA/WT^* animals to *Meox2-Cre* mice (Jackson Laboratory). Once the excision of the STOP cassette was validated by live imaging, *Rosa26^PTPN1-DA/WT^* animals were mated with C57BL/6 mice and compared to *Rosa26^PTPN1-WT/WT^* littermates. Animal procedures described in this study were approved by the McGill University Ethics Committee.

### Pressure overload hypertrophy

Male C57BL/6, PTP1B cKO and PTP1B^F/F^ mice, between 8 and 10 weeks of age, were randomly assigned to groups undergoing sham surgery or constriction of the transverse aorta (TAC) (*65*). Mice were sacrificed 3, 7, 14, or 28 days post-TAC, as indicated. Hearts were harvested, ventricles rapidly removed, snap-frozen in liquid N_2_, and stored at −80°C for protein and RNA extraction.

### PTU treatment

Thyroid hormone deficiency was induced by maintaining animals on an iodine-free chow supplemented with 0.15% propylthiouracil (PTU; TD 97061, Harlan Teklad) for 28 days (*50*).

### Transthoracic echocardiography

Transthoracic echocardiography was performed using an i13L probe (10-14 MHz) linked to Vivid 7 Dimension system (GE Healthcare Ultrasound), with mice being sedated by 1.5-2% isofluorane. Two-dimensional echocardiography was used to measure dimension of left ventricular outflow tract (LVOT), and of ascending aorta proximal to the banding area, banding area, and of descending aorta distal to the banding area. Cross banding peak velocity, peak and mean gradient (V _Peak_, G _Peak_, G _Mean_) were measured by enlarged pulsed wave Doppler. Thickness of LV anterior and posterior wall at end diastole (LVAW_d_, LVPW_d_), LV dimension at end diastole and systole (LVD_d_, LVD_s_), and left atrium dimension at end cardiac systole (LAD_s_) were measured by M-mode echocardiography. LV mass was calculated using appropriate formulae (*66*). LV fractional shortening and ejection fraction (FS, EF) were obtained by formulae available within Vivid 7. Trans mitral flow peak velocity in early (E) and atrial (A) filling, E deceleration time (EDT), deceleration rate (EDR), time interval from mitral valve closure to opening (MV_CO_) were measured by PW; so were velocity of systolic and diastolic (S, D) wave in both left lower and upper pulmonary venous flow (PVF), and S wave deceleration slope in lower PVF. LV ejection time (LVET), stroke volume (SV) and cardiac output (CO) were measured from LVOT flow obtained by PW. LV global myocardial performance index (MPI _Global_) was calculated as (MV_CO_-LVET)/LVET X 100 %. LV iso-volumetric relaxation time (IVRT) was measured by enlarged PW, corrected by R-R interval on simultaneously recorded ECG. Mitral lateral and septal annulus moving velocity in systole (S_L_, S_S_), early and atrial diastole (e’, a’), or fused diastole (f’), and time interval from ending of a’ to beginning of e’(b), from beginning to ending of S_L_ / S_S_ (a) were measured by tissue Doppler imaging. Average of three consecutive cardiac cycles was used for all measurements. The experimenter was blinded to the genotype of the mice.

### Gravimetric analysis

Mice were weighed and executed by cervical vertebra luxation. Hearts were excised, briefly washed with PBS, weighed and snap-frozen in liquid N_2_-cooled isopentane. A Vernier caliper was employed to measure the left tibia length of mice.

### Histology and morphology

Staining and determination of cardiomyocyte area were performed in the histology facility in the laboratory of Dr. Martin Sirois at the Montreal Heart Institute by personnel blinded to the genotype and treatment of mice. Hearts were excised and immediately placed in 10% neutral-buffered formalin at room temperature for 24 h after a brief rinse with PBS. Specimens were embedded in paraffin and sliced into 1 mm sections for hematoxylin and eosin (H&E) staining. Images were taken using an Olympus BX46 microscope. For Texas Red-X conjugated wheat germ agglutinin staining (WGA), hearts were snap-frozen in liquid N_2_-cooled isopentane before cutting into 14 μm sections with a cryostat. The sections were fixed with 4% neutral-buffered formalin, rinsed with PBS then blocked using donkey serum. Sections were stained with Texas Red-X conjugated WGA (Molecular probes) for 60 min and washed with PBS. Cardiomyocyte cross-sectional areas were calculated on a digital microscope using the Image Pro Plus software.

### Adult mouse ventricular cardiomyocyte isolation

Adult mouse ventricular cardiomyocytes were isolated by enzymatic dispersion on a modified Langendorff apparatus as previously described (*67*). Briefly, after cannulation, hearts were rapidly perfused with the following solution A (130 mM NaCl, 15 mM KCl, 0.6 mM KH_2_PO_4_, 0.6 mM Na_2_HPO_4_, 1.2 mM MgSO_4_-7H_2_O, 10 mM HEPES, 4.6 mM NaHCO_3_, 30 mM taurine, 5.5 mM glucose, 5 μM blebbistatin) for 5 min at 3 ml/min. Next, the digestion buffer consisting of 50 ml of solution A supplemented with 120 mg of collagenase type II (290 U/ml, Worthington) was used for enzymatic dispersion. pH was adjusted to 7.4 with NaOH. After 7 to 9 min of digestion, a stopping buffer made from solution A supplemented with 10% FBS and 12.5 M Ca^2+^ was used. The ventricles were cut, minced, and triturated to yield individual rod-shaped myocytes.

### Neonatal rat ventricular cardiomyocyte isolation

Neonatal rat ventricular cardiomyocytes (NRVM) were isolated as previously described (*68*). Briefly, hearts were removed from 2-day old rats killed by decapitation. Ventricular tissues were digested with 0.1% trypsin (Worthington) in HBSS (Wisent laboratories) overnight at 4°C. Ventricular cells were then recovered by repeated digestions of the tissue in 10 ml of 0.1% collagenase in HBSS. The supernatants collected from each digestion were first centrifuged at 1000 rpm for 3 min at 4°C. The pellets were resuspended in ice cold HBSS, pooled and centrifuged at 1000 rpm for 4 min at 4°C. Cells were resuspended in DME containing 7% FBS and preplated in T75 flasks (Corning Inc.) for 75 min twice to enrich for myocytes and decrease contamination by non-muscle cells. Non-adherent cells were then plated at a density of 1000 cells/mm^2^ in 60 mm dishes. After 24 h, the culture medium was changed to serum-free medium containing insulin (5 μg/ml), transferrin (5 pg/ml), and sodium selenite (5 ng/ml) (ITS media). All experiments were performed 24 h later.

### Cell lines and retroviral infection

NRVMs were maintained in DMEM containing serum-free ITS and penicillin/streptomycin whereas HEK293T cells (ATCC) were cultured in DMEM containing 10% FBS, penicillin/streptomycin. Viruses were generated using HEK293T cells as previously described (*69*) and cardiomyocytes were incubated with virus-containing media. Media was replaced 12 h later to prevent cell toxicity. The infected cell populations were lysed for further experiments at the indicated times post-infection.

### Lysate preparation, immunoprecipitation and immunoblotting

At the end of each treatment, hearts were harvested and ventricles were rapidly removed, snap-frozen in liquid N_2_, and stored at −80°C. Ventricles were subsequently pulverized under liquid N_2_ and the powder was resuspended, using a Potter-Elvehjem tissue grinder (10 strokes), in 1 ml of ice-cold lysis buffer (50 mM Tris (pH 7.5), 150 mM NaCl, 50 mM NaF, 2 mM EDTA, 5 mM Na_3_VO_4_, 0.1% SDS, 0.5% v/v deoxycholic acid, 10 μg/ml Rnase A, 5 mM DTT, and 1% v/v NP40 and proteases inhibitors). For immunoprecipitations, 400 μg of protein were incubated with 20 μl of PT-66 anti-phosphotyrosine antibody cross-linked to Protein A-Sepharose (Pharmacia) at 4°C for 3 h as previously described (*31*). Immune complexes were washed 3 times with lysis buffer. Washed immunoprecipitates were eluted by the addition of 20 μl of 4x Laemmli sample buffer. The proteins were separated by electrophoresis on 10% SDS-PAGE, samples were transferred at 100 V and 5 °C for 90 min onto 0.2 μM nitrocellulose membranes in a transfer buffer containing CAPS (10 mM CAPS, 10% methanol, pH 11) to transfer AGO2, or a Tris-glycine transfer buffer (25 mM Tris, 192 mM glycine, 5% methanol) for all other proteins. Membranes were blocked for 1 h in a solution comprising 5% (w/v) skimmed milk powder (Carnation) in TBST (25 mM Tris (pH 7.5), 150 mM NaCl and 0.05% (v/v) Tween-20). Membranes were incubated with primary antibodies in 1% BSA in TBST, for 16 h at 4°C. After washing 3 times with TBST, membranes were reblocked with TBST containing 5% skimmed milk for a 10 min period, and incubated in the presence of HRP-coupled secondary antibodies. Following a final round of washes with TBST, phosphorylated proteins were visualized using ECL according to the manufacturer’s instructions and visualized using BML or BMR films (Kodak), or ChemiDoc MP Imaging System (BioRad).

### Assay of PTP1B oxidation

The cysteinyl-labeling assay of reversibly oxidized PTP1B was performed as previously described with some modifications (*17, 18*). To prevent the spontaneous oxidation of PTP1B by ambient oxygen, the lysis buffer (50 mM Sodium Acetate (pH 5.5), 150 mM NaCl, 10% glycerol, 1% Surfact-Amp NP-40, 10 μg/ml aprotinin, 10 μg/ml leupeptin, 100 U/ml SOD, 100 U/ml catalase, 50 mM IAA,) was degassed and placed on ice into an hypoxic glove-box station equilibrated with 100% argon. Snap-frozen ventricles were moved into the hypoxic glove box station and pulverized under liquid N_2_. The powder was resuspended in 800 μl of ice-cold lysis buffer using a Potter-Elvehjem and samples were transferred in amber-colored Eppendorf tubes and left to alkylate for 1 h on a shaker at room temperature. Following the buffer-exchange step to remove the excess IAA from the lysates using desalting columns, 1 mM TCEP was added to reduce and reactivate reversibly oxidized PTP1B. Following this step, 5 mM sulphydryl-reactive IAP-biotin probe was added to the lysate, to label the reactivated PTP1B. Labelled PTP1B was then pulled-down by streptavidin-Sepharose beads, washed with lysis buffer (pH 5.5) and resuspended in 20 μl 4X Laemmli sample buffer and heated at 90°C for 90 s.

### PTP1B substrate-trapping assay

AGO2 was confirmed as a substrate of PTP1B in H-Ras^V12^-expressing cardiomyocytes and in hearts following a modified protocol previously described by Yang et al. (*31, 45*). RNVMs and hearts from transgenic mice expressing the catalytically impaired PTP1B trapping mutant [PTP1B D181A, PTP1B (DA)] were lysed in a substrate-trapping lysis buffer (50 mM Tris, 5 mM EDTA, 150 mM NaCl, 1% Triton X-100, 5 mM IAA, 10 mM Na_2_HPO_4_, 10 mM NaF), supplemented with 10 μg/ml RNAse A. Lysates were then incubated with anti-Flag beads (Dynabeads) for 90 minutes at 4°C, and the immunoprecipitated proteins were washed with LB. Analysis was performed by western blot.

### RNA isolation and microarray analysis

Hearts were harvested and ventricles were rapidly removed, snap-frozen in liquid N_2_, and stored at −80°C. Ventricles were pulverized under liquid N_2_ and total RNA was extracted using Trizol. Microarray analysis was performed in the Genome Quebec facility at McGill University, using the GeneChip miRNA 4.0 Array (Affimetrix) following manufacturer’s instructions. Differential analysis was performed using LIMMA, a linear model fit to each gene separately. Normalized expression values were reported, together with expression values centered in reference to a specific control group. A heatmap was generated where Euclidean distance was used for hierarchical clustering of miRNA hits.

### AGO2 immunoprecipitation

AGO2 immunoprecipitation was performed as previously described (*31*) with minor modifications. Ventricles were pulverized under liquid N_2_ and the powder was resuspended resuspended in lysis buffer LB AGO (10 mM HEPES pH 7.0, 100 mM KCl, 5 mM MgCl, 0,5% NP-40, 1% Triton X-100, 10% glycerol, 1mM DTT, EDTA free protease inhibitors, RNASE out), homogenized (Dounce homogenizer, 20 passes) and incubated on a clinical rotator at 4°C for 30 min. Lysates were centrifuged at 14,000 X g for 10 min at 4°C. Protein concentration was determined using the method of Bradford (*70*) using γ-globulin as standard. 2 mg of ventricular lysate were incubated with AGO2 antibody (pre-bound to protein A Dynabeads) on a clinical rotator for 4 h at 4°C. The beads were then collected using a magnetic stand and washed twice with NT2 buffer (50 mM Tris (pH 7.4), 150 mM NaCl, 1 mM MgCl_2_, 0.05% NP-40, 100 U/ml RNasin, 1 mM DTT) for 10-15 minutes and two additional times with a higher salt-containing NT2 buffer (300 mM NaCl). Tubes were changed for the last wash. As a control, total RNA extraction was performed with 10% of lysate. RNAs were extracted from both lysates and immunoprecipitates using Trizol.

### Quantitation of AGO2-bound RNAs

qRT-PCR of AGO2-bound mRNAs was performed as described previously (*31*). Briefly, RNAs from immunoprecipitated AGO2 and lysates were extracted using Trizol and treated with RNase free DNase. Following DNase treatment, cDNAs were synthesized using a cDNA synthesis kit. The cDNA library was then used as template for qRT-PCR using a SYBR Green master mix. AGO2-bound miRNA were quantified as described with some modifications (*71*). Briefly miRNA were reverse-transcribed using miRNA-specific stem-loop primers. Following a pre-amplification step to increase the amount of cDNA and to improve the sensitivity of the TaqMan qPCR reaction, quantitation of AGO2-bound miRNAs was performed in a 20 μl volume containing multiplexed AGO2-bound cDNAs, TaqMan Universal PCR Master Mix and TaqMan specific probe with RT-qPCR amplifications conducted as recommended by the manufacturer.

### Real-time quantitative PCR

qRT-PCR was performed to quantify relative transcript levels of hypertrophy markers in our treated and untreated mouse groups. Snap-frozen ventricles were pulverized under liquid N_2_ and total RNA was extracted using Trizol. cDNA was synthesized using M-MLV reverse transcriptase (Invitrogen) using random primers. RT-PCR results from each primer pair were normalized to those of GAPDH (forward: 5’-CTGCACCACCAACTGCTTAGC-3’, reverse: 5’-ACTGTGGTCATGAGCCCTTCCA-3’) gene expression and were compared across conditions. The primers used for fetal cardiac genes, ANP (forward: 5’-GTGCGGTGTCCAACACAGA-3’, reverse: 5’-TTCTACCGGCATCTTCTCCTC-3’), BNP (forward: 5’-GTTTGGGCTGTAACGCACTGA-3’, reverse: 5’-GAAAGAGACCCAGGCAGAGTCA-3’) and β-MHC (forward: 5’-AGGGTGGCAAAGTCACTGCT-3’, reverse: 5’-CATCACCTGGTCCTCCTTCA-3’). All RT-qPCRs were carried out in duplicate and were performed on the MX3000p thermal cycler system from Stratagene with the following conditions: one denaturing step at 95°C for 10min, followed by 40 cycles consisting of denaturing at 95°C for 15 s and annealing and elongation at 60°C for 60 s, followed by an inactivation step of 10 min at 99.9°C.

### Statistics

Data are expressed as the mean ± SEM. Testing differences in means between 2 groups was done by unpaired t tests, whereas those among more than 2 groups were by ANOVA followed by a post-hoc Bonferroni’s comparison test. *P* < 0.05 was considered to be statistically significant.

## SUPPLEMENTARY MATERIALS

Supplementary Materials and Methods

Table S1. Echocardiography of PTP1B^F/F^ and PTP1B cKO Mice

Table S2. Echocardiography of PTP1B^F/F^ and PTP1B cKO mice after TAC

Table S3. Echocardiography of PTP1B^F/F^ and PTP1B cKO mice after TAC and PTU

Figure S1. Changes in miRNA expression in TAC-treated PTP1B cKO and PTP1B^F/F^ mice.

Figure S2. Generation of PTP1B (D181A) trapping-mutant transgenic mice.

Figure S3. AGO2 phosphorylation in hearts following TAC.

Figure S4. Left ventricular filling patterns of sham and TAC hearts from PTU-treated mice.

Figure S5. Quantification of α-MHC and β-MHC mRNA levels in PTU-treated mice.

Figure S6. Changes in relative MHC isoform expression ratio in TAC and PTU-treated mice.

## Acknowledgements

We thank Dr. Linda Van Aelst for the retroviral vector pBabe–H-RASV12-Puro and Dr. Nicholas K. Tonks for the pWZL-PTP1B DA-Hygro expression constructs. We thank Marc-Antoine Gillis, Louis Villeneuve and Noriko Uetani for their technical expertise, Terence Hébert, Bruce Allen, Markus Dagnell, Michael Kapiloff and the members of the Boivin laboratory for helpful discussions.

## Funding

This research was supported by NIH grant HL138605 and an American Heart Association grant 17GRNT33700265 to BB and by Canadian Institutes of Health Research grant MOP-62887 to MLT. BB is also grateful for support from the following foundations: Heart and Stroke Foundation of Canada, SUNY Research Foundation, The Montreal Heart Institute Foundation, Fond de Recherche du Quebec en Santé (FRQS). BB is a FRQS Research Scholar. DPL is a Lewis Katz Young Investigator of the Prostate Cancer Foundation, and is the recipient of a Scholarship for the Next Generation of Scientists from the Cancer Research Society. AB was the recipient of scholarship from the FRQS.

## Author Contributions

G.C. and B.B. designed the experiments. G.C. contributed to all aspects of experimental work. Y.S. and J.C.T. contributed to echocardiography acquisition and provided insightful thoughts and discussion. D.P.L., F.S., A.B., V.V., G.K., B.G.A. and M.L.T. provided critical reagents and contributed to data acquisition. All of the authors contributed to the analysis of the data. G.C. and B.B. wrote the paper. B.B. directed the study.

## Competing Interests

The authors have declared that no conflict of interest exists.

