## Supplemental figures for "Protein Tyrosine Phosphatase 1B Regulates MicroRNA-208b-Argonaute 2 Association and Thyroid Hormone Responsiveness in Cardiac Hypertrophy"

**This file includes:**

Supplementary Material and Methods

Tables S1-S3

Figs. S1-S6

#### SUPPLEMENTARY MATERIAL AND METHODS

##### Reagents and antibodies

PTP1B and THRAP1 antibodies were purchased from Abcam. AGO2 antibodies were purchased from Cell Signaling Technology. H-RAS and anti-phosphotyrosine (4G10) antibodies were purchased from Millipore. Anti-Flag antibody, Dynabeads protein A and protein G were from Thermo Scientific. The β-actin and anti-phosphotyrosine PT-66 beads were from Sigma. HRP-conjugated secondary antibodies were from Jackson Laboratories, GAPDH antibodies and Protein A/G Plus agarose beads were from Santa Cruz Biotechnology. Anti-AGO2 phospho-Tyr$^{393}$ antibodies were generated against N$^{287}$-T-D-P**Y-p**V-R-E-F-G-I-M$^{400}$ and affinity purified at the MRC Protein Phosphorylation and Ubiquitylation Unit (Dundee). Streptavidin-Sepharose beads were purchased from GE Healthcare. Protease inhibitor cocktail tablets and DNase were from Roche. Trizol, cDNA synthesis kit, SYBR Green master mix and TaqMan were from Thermo Scientific. Catalase and superoxide dismutase were from Calbiochem. Surfact-Amps Nonidet P-40, Zeba desalt spin columns, EZ-Link iodoacetyl-PEG2-biotin (IAP-biotin) and iodoacetic acid (IAA) were from Thermo Scientific. RNasin was from Promega. The retroviral vectors pBabe–H-RASV12-Puro and pWZL-PTP1B DA-Hygro were generous gifts from Dr. Linda Van Aelst (*72*) and Dr. Nicholas K. Tonks (*31*), respectively.

##### Live imaging

Mice were anesthetized with 2-chloro-2-(difluoromethoxy)-1,1,1-trifluoro-ethane (isoflurane), injected intraperitoneally with 50 μL luciferin (Caliper Life Sciences), and imaged with the use of the IVIS Spectrum preclinical *in vivo* imaging system (Perkin Elmer), according to the manufacturer's instructions.

#### Table S1. Echocardiography of PTP1B$^{F/F}$ and PTP1B cKO Mice

| | baseline | |
|---|---|---|
| | **F/F mice (n = 17)** | ***PTP1B cKO* (n = 23)** |
| **BW (g)** | 25.6 ± 1.8 | 25.7 ± 2.9 |
| **HR (bpm)** | 483 ± 77 | 411 ± 62* |
| **LVDd (mm)** | 4.09 ± 0.19 | 4.34 ± 0.45* |
| **LVDs (mm)** | 2.68 ± 0.27 | 3.08 ± 0.63* |
| **LVAWd (mm)** | 0.73 ± 0.02 | 0.73 ± 0.06 |
| **LVPWd (mm)** | 0.72 ± 0.04 | 0.72 ± 0.04 |
| **LVM (mg)** | 108.0 ± 10.2 | 119.0 ± 22.2* |
| **LVM/BW (mg/g)** | 4.23 ± 0.34 | 4.64 ± 0.79* |
| **FS, %** | 34.7 ± 4.8 | 29.4 ± 7.2* |
| **EF, %** | 70.3 ± 6.0 | 62.3 ± 11.9* |
| **LAD (mm)** | 2.52 ± 0.16 | 2.45 ± 0.63 |
| **IVRT (msec)** | 5.3 ± 1.4 | 7.1 ± 3.0* |

Two dimensionally guided M-mode echocardiography was performed on 4-week old PTP1B$^{F/F}$ and PTP1B cKO mice. BW, body weight; HR, heart rate; LVDd, left ventricular dimension at end diastole; LVDs, left ventricular dimension at end systole; LVAWd, thickness of left ventricular anterior wall at end diastole; LVPWd, thickness of left ventricular posterior wall at end diastole; LVM, left ventricular mass, FS, left ventricular fractional shortening; EF, left ventricular ejection fraction; LAD, left ventricular internal septum diastolic; IVRT, left ventricular iso-volumetric relaxation time. * $P < 0.05$

#### Table S2. Echocardiography of PTP1B$^{F/F}$ and PTP1B cKO mice after TAC

| | 1m post Sham | | 1m post TAC | |
|---|---|---|---|---|
| | **F/F mice (n = 10)** | ***PTP1B cKO* (n = 14)** | **F/F mice (n = 13)** | ***PTP1B cKO* (n = 17)** |
| **BW (g)** | 28.6 ± 2.7 | 26.6 ± 3.6 | 28.3 ± 3.3 | 28.8 ± 2.8 |
| **HR (bpm)** | 478 ± 40 | 447 ± 61 | 536 ± 48* | 522 ± 66* |
| **LVDd (mm)** | 4.16 ± 0.28 | 4.21 ± 0.28 | 4.19 ± 0.22 | 4.89 ± 0.50*# |
| **LVDs (mm)** | 2.69 ± 0.34 | 2.8 ± 0.19 | 2.91 ± 0.24 | 4.14 ± 0.8*# |
| **LVAWd (mm)** | 0.77 ± 0.04 | 0.75 ± 0.03 | 0.92 ± 0.06* | 0.90 ± 0.07* |
| **LVPWd (mm)** | 0.71 ± 0.04 | 0.74 ± 0.02 | 0.89 ± 0.06* | 0.87 ± 0.1* |
| **LVM (mg)** | 114.1 ± 19.9 | 117.3 ± 15.4 | 150.5 ± 16.4* | 189.0 ± 26.6*# |
| **LVM/BW (mg/g)** | 3.99 ± 0.56 | 4.42 ± 0.28 | 5.38 ± 0.76* | 6.59 ± 0.91*# |
| **FS, %** | 35.4 ± 5.1 | 33.4 ± 4.0 | 30.5 ± 4.3 | 17.9 ± 9.3*# |
| **EF, %** | 71.2 ± 6.2 | 68.7 ± 5.3 | 64.5 ± 6.2* | 41.4 ± 16.1*# |
| **LAD (mm)** | 2.61 ± 0.14 | 2.43 ± 0.20 | 2.62 ± 0.24 | 3.17 ± 0.41*# |
| **IVRT (msec)** | 5.07 ± 0.92 | 5.7 ± 1.62 | 4.81 ± 0.98 | 5.96 ± 2.07# |

Two dimensionally guided M-mode echocardiography was performed 1 month after sham or TAC surgery in PTP1B$^{F/F}$ and PTP1B cKO mice. BW, body weight; HR, heart rate; LVDd, left ventricular dimension at end diastole; LVDs, left ventricular dimension at end systole; LVAWd, thickness of left ventricular anterior wall at end diastole; LVPWd, thickness of left ventricular posterior wall at end diastole; LVM, left ventricular mass, FS, left ventricular fractional shortening; EF, left ventricular ejection fraction; LAD, left ventricular internal septum diastolic; IVRT, left ventricular iso-volumetric relaxation time. * $P < 0.05$ Vs corresponding sham, # $P < 0.05$ vs TAC F/F

### Table S3. Echocardiography of PTP1B$^{F/F}$ and PTP1B cKO mice after TAC and PTU

| | 1m post SHAM | | 1m post TAC | | 1m post SHAM+PTU | | 1m post TAC+PTU | |
|---|---|---|---|---|---|---|---|---|
| | F/F mice (n = 8) | *PTP1b cKO* (n = 10) | F/F mice (n = 9) | *PTP1b cKO* (n = 11) | F/F mice (n = 8) | *PTP1b cKO* (n = 6) | F/F mice (n = 5) | *PTP1b cKO* (n = 10) |
| **BW (g)** | 27 ± 2.27 | 26.0 ± 3.1 | 28.3 ± 3.8 | 28.1 ± 2.9 | 22.3 ± 2.9 | 23.8 ± 2.4 | 22.8 ± 2.9 | 20.3 ± 1.9* |
| **HR (bpm)** | 477 ± 48 | 408 ± 94° | 541 ± 49* | 529 ± 64 | 252 ± 12 | 283 ± 65 | 312 ± 63 | 318 ± 65 |
| **LVDd (mm)** | 4.18 ± 0.35 | 4.09 ± 0.34 | 4.21 ± 0.28 | 5.00 ± 0.43*# | 3.96 ± 0.14 | 3.83 ± 0.26 | 3.99 ± 0.23 | 3.89 ± 0.34 |
| **LVDs (mm)** | 2.74 ± 0.43 | 2.80 ± 0.28 | 2.92 ± 0.22 | 4.14 ± 0.8*# | 2.89 ± 0.22 | 2.79 ± 0.31 | 2.73 ± 0.38 | 2.84 ± 0.41 |
| **LVAWd (mm)** | 0.77 ± 0.05 | 0.79 ± 0.04 | 0.92 ± 0.08* | 0.91 ± 0.07* | 0.69 ± 0.08 | 0.73 ± 0.1 | 0.76 ± 0.1 | 0.83 ± 0.13 |
| **LVPWd (mm)** | 0.72 ± 0.04 | 0.74 ± 0.06 | 0.89 ± 0.19* | 0.89 ± 0.1* | 0.60 ± 0.08 | 0.65 ± 0.09 | 0.73 ± 0.07* | 0.75 ± 0.15 |
| **LVM (mg)** | 116 ± 23.6 | 115.1 ± 13.1 | 153 ± 28.06* | 201.6 ± 45.1*# | 87.4 ± 17.4 | 90.4 ± 17.1 | 107.2 ± 13.5* | 113.2 ± 32.2 |
| **LVM/BW (mg/g)** | 4.28 ± 0.66 | 4.51 ± 0.28 | 5.46 ± 1.02* | 7.26 ± 1.76*# | 3.97 ± 0.74 | 3.84 ± 0.63 | 4.71 ± 0.19 | 5.51 ± 1.22* |
| **FS, %** | 36.6 ± 5.2 | 31.4 ± 5.4 | 30.8 ± 4 | 17.9 ± 9.3*# | 27.1 ± 4.3 | 27.4 ± 4.2 | 32.0 ± 6.5 | 27.2 ± 5.7 |
| **EF, %** | 72.1 ±7.1 | 65.6 ± 8 | 64.9 ± 5.8 | 41.4 ± 16.1*# | 59.3 ± 6.3 | 59.8 ± 6.9 | 66.2 ± 8.6 | 59.2 ± 9.8 |
| **LAD (mm)** | 2.48 ± 0.28 | 2.18 ± 0.18 | 2.64 ± 0.23 | 3.17 ± 0.41*# | 2.29 ± 0.19 | 2.18 ± 0.23 | 2.43 ± 0.47 | 2.09 ± 0.35 |
| **IVRTc** | 0.44 ± 0.06 | 0.44 ± 0.11 | 0.51 ± 0.12 | 0.60 ± 0.21 | 1.24 ± 0.18 | 1.06 ± 0.14 | 1.03 ± 0.03 | 1.08 ± 0.25 |

Two dimensionally guided M-mode echocardiography was performed on sham and TAC PTP1B$^{F/F}$ and PTP1B cKO mice 4 weeks post-surgery, treated or not with PTU . BW, body weight; HR, heart rate; LVDd, left ventricular dimension at end diastole; LVDs, left ventricular dimension at end systole; LVAWd, thickness of left ventricular anterior wall at end diastole; LVPWd, thickness of left ventricular posterior wall at end diastole; LVM, left ventricular mass, FS, left ventricular fractional shortening; EF, left ventricular ejection fraction; LAD, left ventricular internal septum diastolic; IVRTc, heart-rate corrected left ventricular iso-volumetric relaxation time. * $P < 0.05$ Vs corresponding sham, # $P < 0.05$ vs TAC F/F.

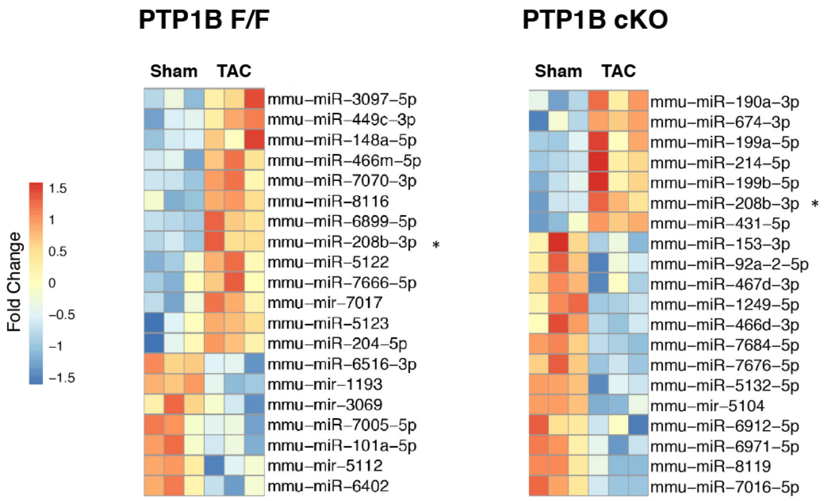


**Figure S1. Changes in miRNA expression in TAC-treated PTP1BcKO and PTP1B$^{F/F}$ mice.** Cardiac miRNA populations were extracted from sham and TAC PTP1B$^{F/F}$ and PTP1B cKO mice 4 weeks post-surgery as described in *Methods*. The RNAs were then analyzed on an Affimetrix 4.0 miRNA Array. Intensity values were transformed into $\log_2$ scale, and the heat map shows $\log_2$ fold change values for each data set (i.e. PTP1B$^{F/F}$ and PTP1B cKO), with red representing relative increased expression levels between sham-operated and 4-week treated hearts and blue representing the relative decrease. The top 20 miRNAs showing the most change in relative $\log_2$ fold change expression are visualized in a heat map generated by Galaxy. miR-208b is identified by *.

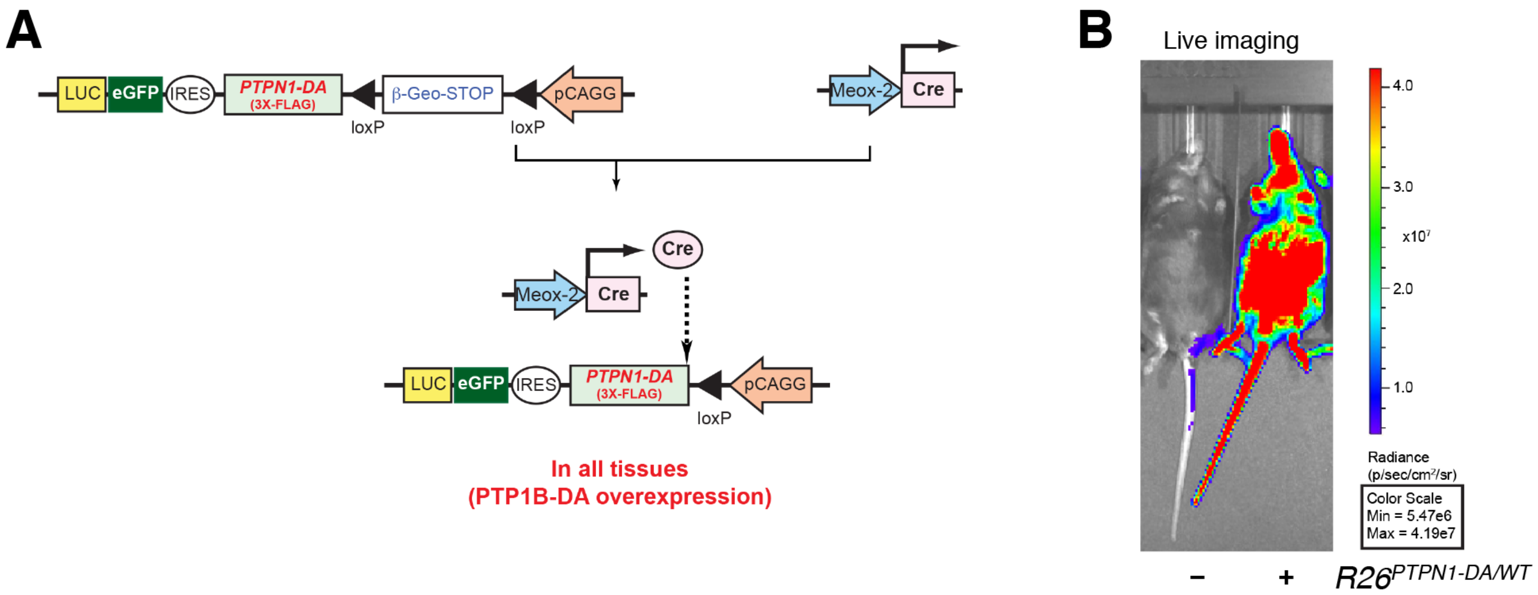


**Figure S2. Generation of PTP1B (D181A) trapping-mutant transgenic mice.** A) *Rosa26*-targeted (*R26*) pCAGG-promoter-based construct drives strong expression of 3XFLAG-PTP1B protein (*R26PTPN1/WT*), together with an eGFP/luciferase reporter in the whole body, following breeding with a *Meox2-Cre* mouse. B) Validation of the $R26^{PTPN1/WT}$ transgene by live imaging.

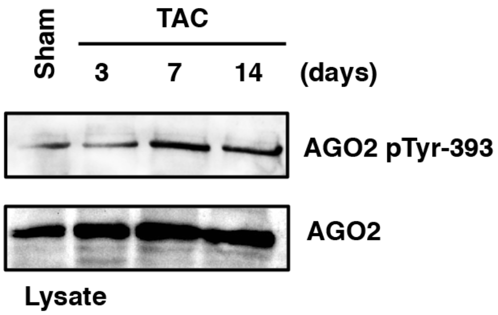


**Figure S3. AGO2 phosphorylation in hearts following TAC.** Lysates from sham-operated hearts or from hearts subjected to 3, 7 or 14 days of TAC-induced pressure overload were prepared as described in *Methods*, resolved by SDS-PAGE, transferred to nitrocellulose, and probed for AGO2 phospho-$Tyr^{393}$ and total AGO2 immunoreactivity.

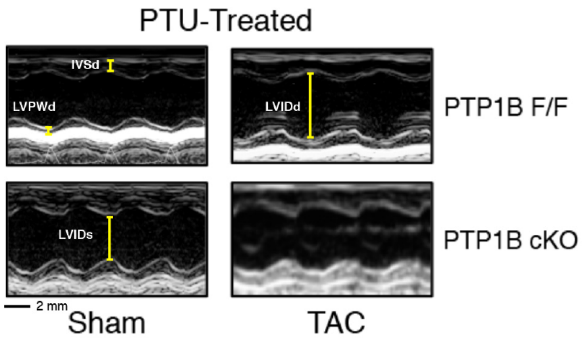


**Figure S4. Left ventricular filling patterns of sham and TAC hearts from PTU-treated mice.** Representative M-mode echocardiograms of sham and TAC PTP1B$^{F/F}$ and PTP1B cKO mice 4 weeks post-surgery, fed with or without PTU-containing chow. The IVSd (interventricular septal end diastole) and LVPWd (left ventricular posterior wall end diastole) dimensions are used to determine left ventricular hypertrophy, LVIDd and LVIDs (left ventricular internal diameter end diastole and end systole, respectively) are used to calculate fractional shortening (FS).

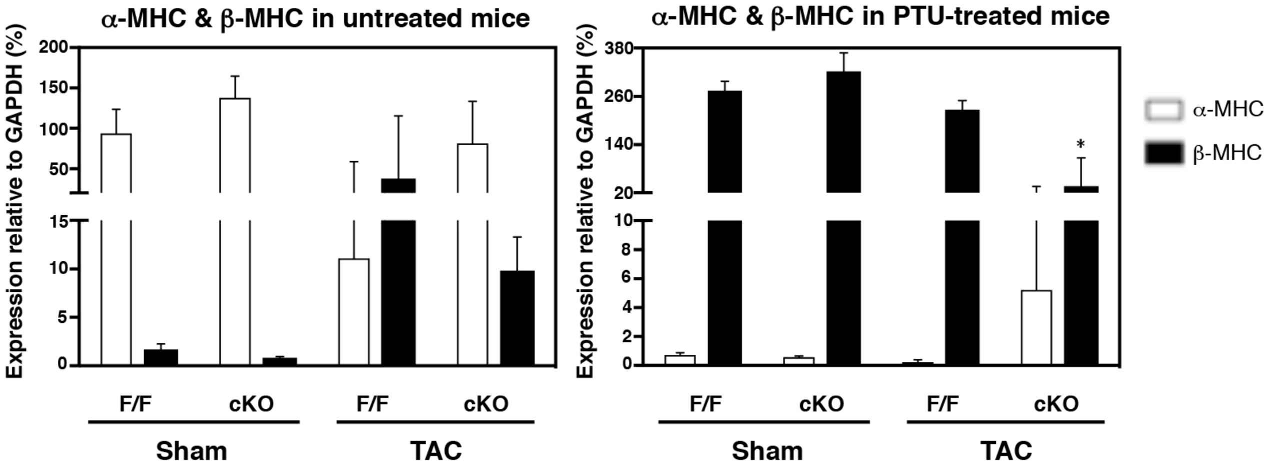


**Figure S5. Quantification of α-MHC and β-MHC mRNA levels in PTU-treated mice.** Relative expression of α-MHC and β-MHC in hearts from sham and TAC PTP1B$^{F/F}$ and PTP1B cKO mice 4 weeks post-surgery, treated or not with PTU. Expression levels of mRNAs are normalized to GAPDH. Representative data are presented as mean ± SEM for three or more independent experiments. Statistical analyses were done by two-way ANOVA with Bonferonni's post hoc test for comparisons among multiple groups. * $P < 0.01$ vs. respective sham-operated groups.

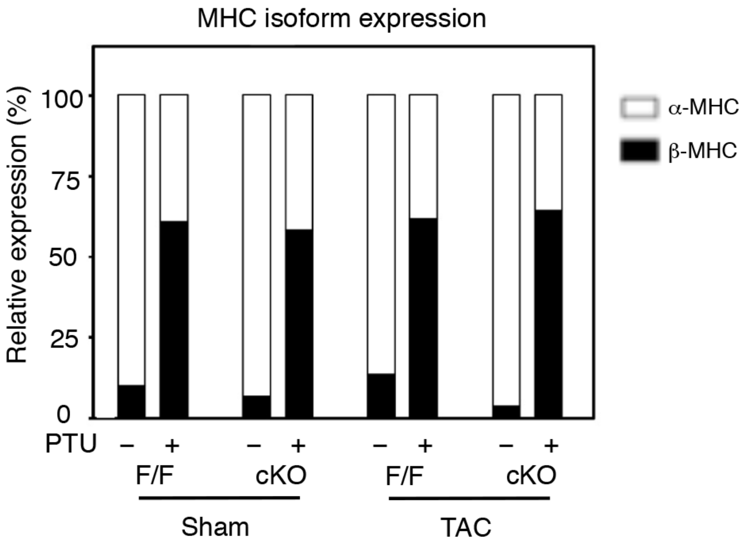


**Figure S6. Changes in relative MHC isoform expression ratio in TAC and PTU-treated mice.** Relative expression of α-MHC and β-MHC in sham and TAC hearts from PTP1B$^{F/F}$ and PTP1B cKO mice treated or not with PTU for 4 weeks. Expression data is normalized to the established 90:10 expression ratio for α-MHC:β-MHC isoforms in mice, as previously reported (*38*).
